# Molecular logic of neuromodulatory systems in the zebrafish telencephalon

**DOI:** 10.1101/2023.10.04.560846

**Authors:** Lukas Anneser, Chie Satou, Hans-Rudolf Hotz, Rainer W. Friedrich

**Author notes:** Corresponding author: Dr. Rainer Friedrich.

## Abstract

The function of neuronal networks is determined not only by synaptic connectivity but also by neuromodulatory systems that broadcast information via distributed connections and volume transmission. To understand the molecular constraints that organize neuromodulatory signaling in the telencephalon of adult zebrafish we used transcriptomics and additional approaches to delineate cell types, to determine their phylogenetic conservation, and to map the expression of marker genes at high granularity. The combinatorial expression of GPCRs and cell type markers indicates that all neuronal cell types are subject to modulation by multiple monoaminergic systems and distinct combinations of neuropeptides. Individual cell types were associated with multiple (typically >30) neuromodulatory signaling networks but expressed only a few diagnostic GPCRs at high levels, suggesting that different neuromodulatory systems act in combination albeit with unequal weights. These results provide a detailed map of cell types and brain areas in the zebrafish telencephalon, identify core components of neuromodulatory networks, highlight the cell type-specificity of neuropeptides and GPCRs, and begin to decipher the logic of combinatorial neuromodulation.

## Introduction

Neuronal circuit architecture is characterized by distinct cell types that interact in specific configurations. Information flow within these circuits depends on the synaptic connectivity between individual neurons, which is currently analyzed by the reconstruction of synaptic wiring diagrams using volume electron microscopy techniques^1,2^. In addition, information is transmitted by neuromodulatory systems through distributed connectivity and volume transmission. This additional layer of signaling exists across the entire animal kingdom and exhibits substantial complexity even in small circuits^3,4^. Progress over the last decades revealed that individual cell types are typically influenced by multiple modulatory systems, indicating that neuromodulation follows a combinatorial logic. A full understanding of neural circuits should therefore include a comprehensive analysis of neuromodulatory systems and their interactions at the level of transcriptionally defined cell types^5,6^.

Here we analyze neuromodulatory networks in the zebrafish telencephalon using next-generation sequencing approaches. Owing to its small size, the zebrafish offers outstanding opportunities to study mechanisms of cognition in a vertebrate by combining optical imaging, circuit reconstruction, and molecular approaches^7^. The telencephalon is of particular interest because it plays important roles in higher brain functions such as prediction^8^, social behavior^9,10^, and associative learning^11^.

The organization of the zebrafish brain has been described based on anatomical^12,13^, neurodevelopmental^14^, genetic^15^, and functional^16^ approaches. Comparisons to other vertebrate classes are complicated by the fact that the teleostean telencephalon develops by a morphogenetic process termed eversion that differs from evagination in amniotes^14^. Gene expression studies are therefore a promising approach to determine phylogenetic relationships between cell types and brain areas of teleosts and other vertebrate classes. Homologies have been established for a subset of telencephalic areas in teleosts such as the posterior zone of the dorsal pallium (Dp), which most likely corresponds to piriform cortex in the lateral pallium of amniotes^13^. However, potential homologies of various other neuroanatomical areas have been proposed but not firmly established.

Information processing is influenced throughout the vertebrate telencephalon by classical neuromodulatory systems such as the dopaminergic, serotonergic, or cholinergic systems, often within the same circuit^17,18^. While previous studies focused primarily on cellular effects and functional roles of individual neuromodulators, neuronal circuit function is most likely controlled by multiple systems acting in concert. This combinatorial view of neuromodulation has emerged from pioneering studies in invertebrates^3,5,19^ but most likely also applies to the vertebrate brain where different neuromodulators can be co-released^17^ and jointly regulate network function^18,20^. Computationally, the co-release of neurotransmitters and one or multiple neuromodulators can increase the degrees of freedom for individual neurons in a circuit by modulating pre- and postsynaptic components and adding different temporal scales of action^19^. A systematic analysis of combinatorial neuromodulation would benefit greatly from comprehensive information about the co-expression of multiple neuromodulators and their receptors in individual neurons and cell types. Such joint gene expression maps would provide a basis to infer whether specific cell types are potential senders or receivers of distinct neuromodulatory signals^6^, to detect correlated effects of neuromodulation on different cell types, and to explore interactions between different neuromodulatory systems or subsystems.

We used transcriptomic methods to analyze combinatorial gene expression in individual neurons. Previously, transcriptomic approaches showed that neuropeptide precursors are often among the most highly expressed genes in neurons^21,22^. Moreover, transcriptomic analyses revealed that the combinatorial expression of neuropeptides and their cognate receptors is cell type-specific and a conserved feature of cortical circuits^6,22^. We extended these approaches and focused specifically on G protein-coupled receptors (GPCRs) for neuropeptides and classical monoaminergic neuromodulators.

Our results identify neuromodulatory signatures for all major pallial areas and neuronal cell types in the telencephalon. We localized individual GABAergic and glutamatergic cell types within their anatomical context and provide evidence of evolutionary conservation. All neuronal cell types were found to be embedded in multiple neuromodulatory networks. This comprehensive analysis provides new insights into the organization of cell types and neuromodulatory systems in the telencephalon and may serve as a basis for further functional analyses of neuromodulatory networks in cognition and behavior^23–25^.

## Results

### Pallial regions have distinct modulatory signatures

The potential impact of a neuromodulator on a neuronal circuit is constrained by the receptors expressed in the constituent neurons. Our first aim was to comprehensively analyze the expression of neuromodulatory receptors in the major pallial subregions of the zebrafish telencephalon, the central (Dc), medial (Dm), lateral (Dl), and posterior (Dp) zones of the dorsal pallium^26^ (Fig. 1A). We dissected these pallial areas from adult zebrafish brains and performed bulk RNA sequencing (Fig. S1A). Gene expression patterns from different brain areas were separated by projections onto the first few principal components (PCs; Fig. 1B; Fig. S1B), implying region-specific differences in gene expression profiles. Indeed, we identified 7,164 genes that were differentially expressed between regions. We then mined the KEGG database for G protein-coupled receptors (GPCRs, search term dre04030) and found 270 GPCRs to be expressed in the telencephalon. Out of these, 150 were part of known neuromodulatory systems, as determined by manual curation, and 59 out of these 150 GPCRs (39.3 %) were differentially expressed across brain areas (Fig. 1C; see Methods for criteria). To determine whether the numbers of differentially expressed GPCRs were equally distributed among pallial areas, we averaged expression patterns across replicates and visualized the result in a heatmap (Fig. 1D). Differentially expressed GPCRs were found most frequently in Dc or Dm but locally enriched subsets of GPCRs were found in all pallial regions (Fig. 1D). Anatomically defined pallial brain areas therefore exhibit characteristic signatures of GPCR expression.

**Figure 1.**
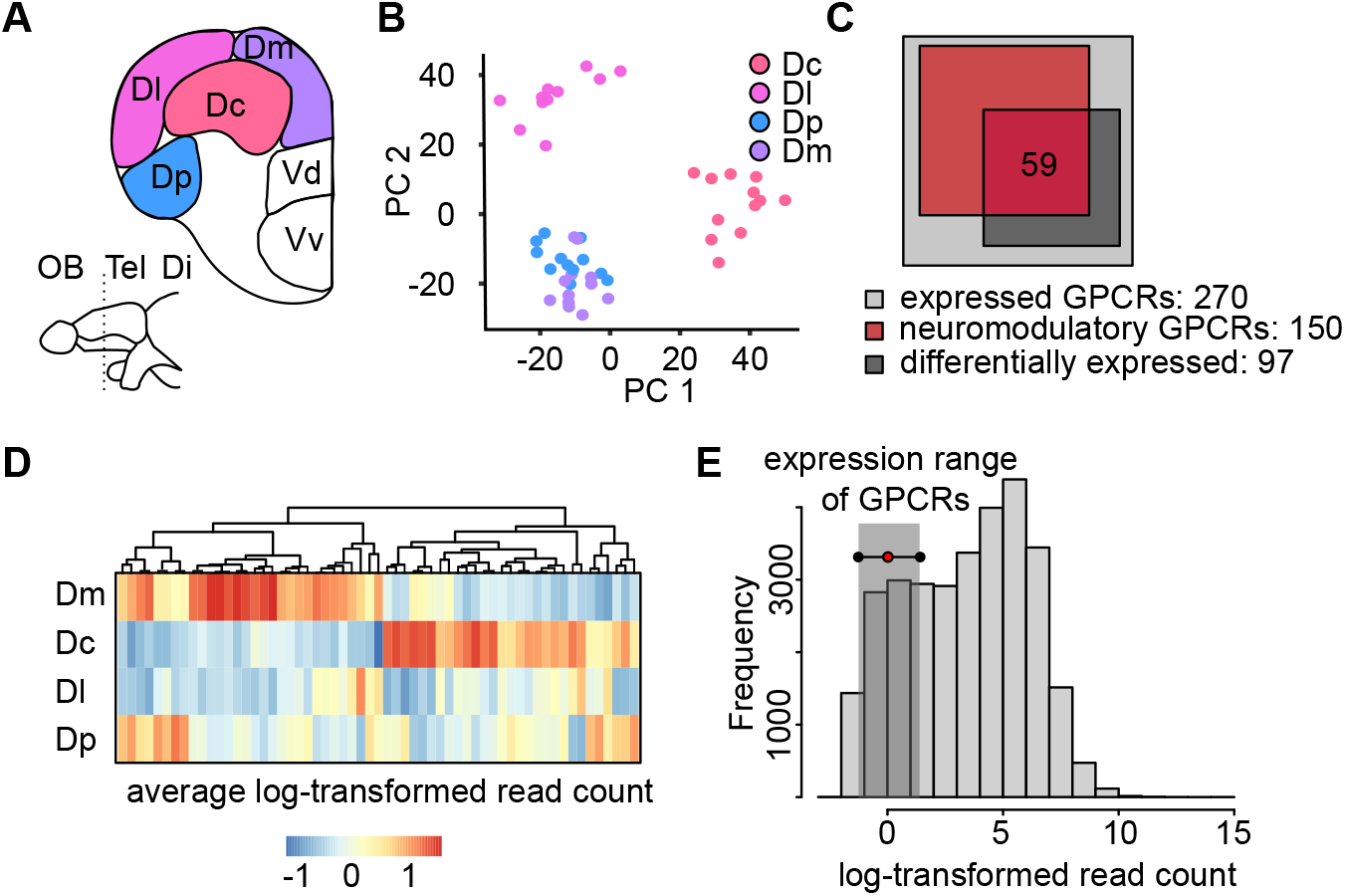
Transcriptomic profile of pallial regions based on bulk sequencing. (**A**) Coronal section through the adult zebrafish telencephalon showing the four pallial regions (Dc, Dm, Dl, Dp) dissected for bulk sequencing. Tel, telencephalon; Di, diencephalon; OB, olfactory bulb. (**B**) Expression patterns of the 2.000 most variable genes projected onto the first two PCs. Each data point is one replicate from a pallial region (color-coded). Dp and Dm can be separated in projections onto PCs 3 and 4 (Fig S1B). (**C**) Venn diagram showing the number of GPCRs that were differentially expressed across brain areas and associated with neuromodulatory systems. Expression analysis was performed using the intersection between these sets (n = 59 genes). (**D**) Expression of the 59 GPCRs across pallial brain regions, sorted by clustering. (**E**) Histogram showing expression levels of different genes, quantified by averaged read count, and expression range of GPCRs.

### Identification of cell types in the telencephalon

As data obtained from bulk sequencing conflates molecular signatures of different cell types, we used single-cell transcriptomics to obtain a more granular understanding of GPCR expression. We dissected the telencephalon of adult zebrafish, removed the olfactory bulbs to increase coverage of pallial and subpallial regions, and performed single-cell RNA-sequencing using the 10X Genomics approach (Fig. 1A). Because GPCRs are expressed at comparatively low levels (Fig. 1E) we adjusted sequencing parameters to reach approximately 8000 unique molecular identifiers per cell^23^. This approach resulted in reads on 2753 genes on average, substantially more than in previous studies using zebrafish (Fig. S2A-C). After quality control, we obtained a total number of 16,321 telencephalic cells that showed a highly clustered distribution in UMAP space (Fig. 2A). Moreover, cells could be clearly assigned to superclusters that were previously described in similar approaches across species (Fig. 2B, C, S2D-F)^27,28^. For example, neurons could be identified by the expression of *snap25a*^29^ and further subdivided into GABAergic and glutamatergic cells using the marker genes *slc32a1* and *slc17a6a*^30,31^. Ependymoglial cells were identified by expression of *gfap*^32^, oligodendrocyte precursor cells (OPCs) by *olig1*^33^, differentiating oligodendrocytes by *gpr17*^34^, and mature oligodendrocytes by *mbpa*^35^, all consistent with previous results from zebrafish^28,36,37^ and other species^27,38,39^. We also found a few cells with characteristic signatures of habenular neurons (*kiss1^+^*/*gng8^+^*), presumably because the dissected tissue contained parts of the habenula.

**Figure 2.**
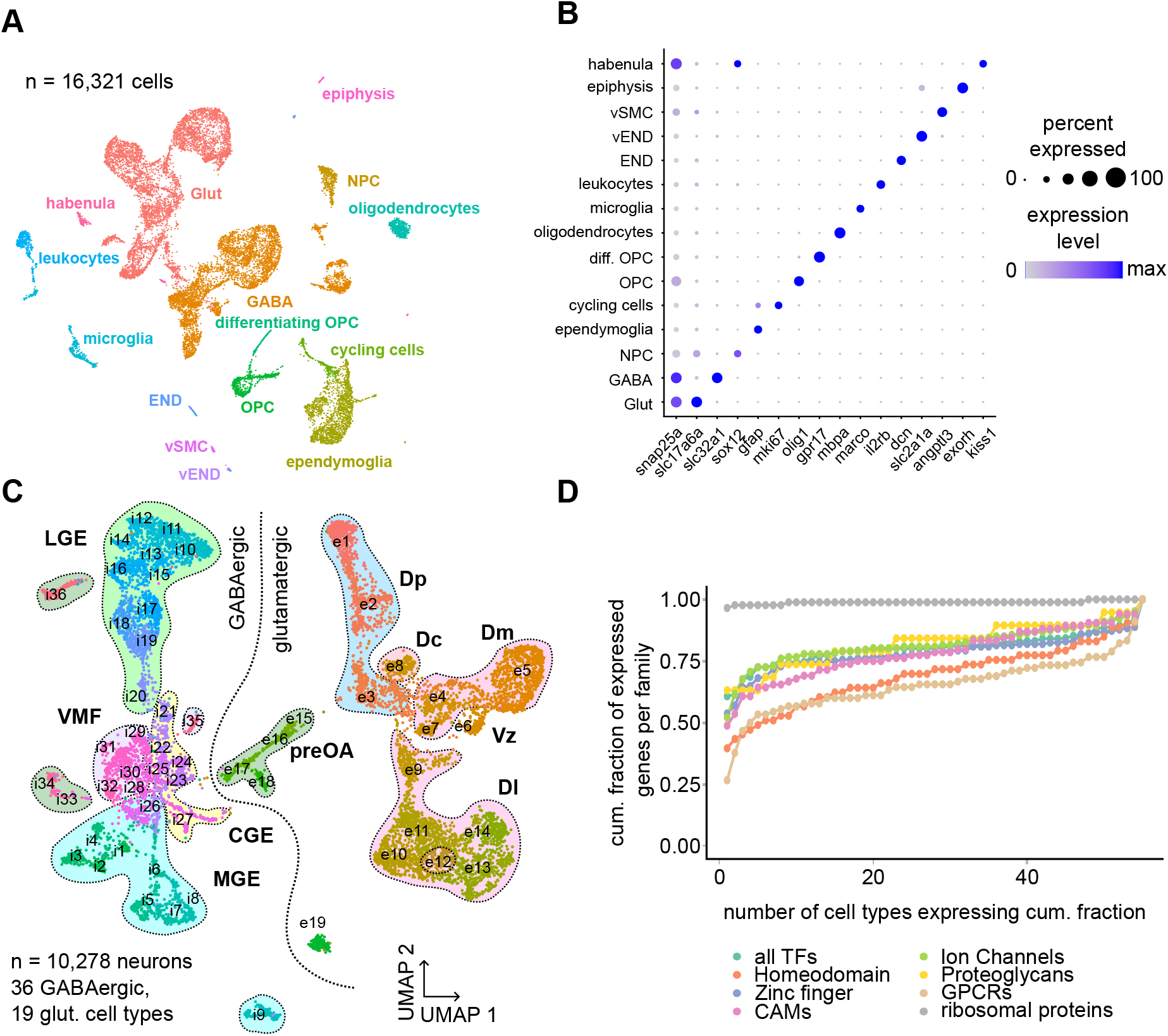
scRNA-seq of the adult zebrafish telencephalon reveals neuronal diversity. (**A**) Uniform manifold approximation and projection (UMAP) representation of the 16,321 sequenced cells, color-coded by cell type clustering. END, endothelial cells; vEND, vascular endothelial cells; vSMC, vascular smooth muscle cells; OPC, oligodendrocyte precursor cells; NPC; neural progenitor cells. (**B**) Expression of selected marker genes in individual cell types. Color indicates expression level (normalized log read count), and dot size represents fraction of expressing cells per cluster. (**C**) UMAP plot of 10,278 neuronal cell types clustered separately. Association of cell types with brain area is indicated by shaded areas. LGE, lateral ganglionic eminence; VMF, ventromedial forebrain; CGE, caudal ganglionic eminence; MGE, medial ganglionic eminence; preOA, preoptic area; Dp, posterior zone of the dorsal pallium; Dc, central zone of the dorsal pallium; Dm, medial zone of the dorsal pallium; Dl, lateral zone of the dorsal pallium. (**D**) Cumulative fraction of cell types expressing different gene families (TFs, transcription factors; Homeodomain TFs; Zinc finger TFs, CAMs, cell adhesion molecules; ion channels; proteoglycans; ribosomal proteins; GPCRs).

We then focused on the 4,733 GABAergic and 5,545 glutamatergic cells (excluding habenular neurons), which we separately subclustered for a more fine-grained analysis of neuronal cell type diversity. Similar approaches in reptiles and amphibians identified 26 - 47 glutamatergic and 18 - 67 GABAergic cell types in the telencephalon^27,38–40^. In our dataset we found 19 glutamatergic and 36 GABAergic putative cell types (Fig. 2C, S2G-I).

Cell classes can be defined by the combinatorial expression of relevant transcription factors (TFs), which jointly select the genetic programs characterizing the cell^41^. As transcription-based cell type taxonomy is a hierarchical approach ranging from stringently defined superclusters to more fuzzy but fine-grained cell types^42^ we examined whether neuronal cell types can also be identified by a purely TF-based classification. We found that TFs were more selectively expressed across cell types than other gene families, which was particularly pronounced for homeodomain TFs (Fig. 2D, 3A, S3A). We then trained a random forest classifier to assign cell type identity based on the expression of TFs. This classifier successfully retrieved almost all 55 neuronal cell types that were previously identified based on the expression matrix containing all genes (Fig. S3B, C) and allowed us to retrieve genes that were particularly informative about cell type assignment (Fig. S3F-H). Using this purely data-driven approach, we identified multiple TFs commonly used in cell type classification in the telencephalon, such as *nr2f2*, *zic1*, and *foxg1a* (Fig. 3A, S3F)^27,40^.

**Figure 3.**
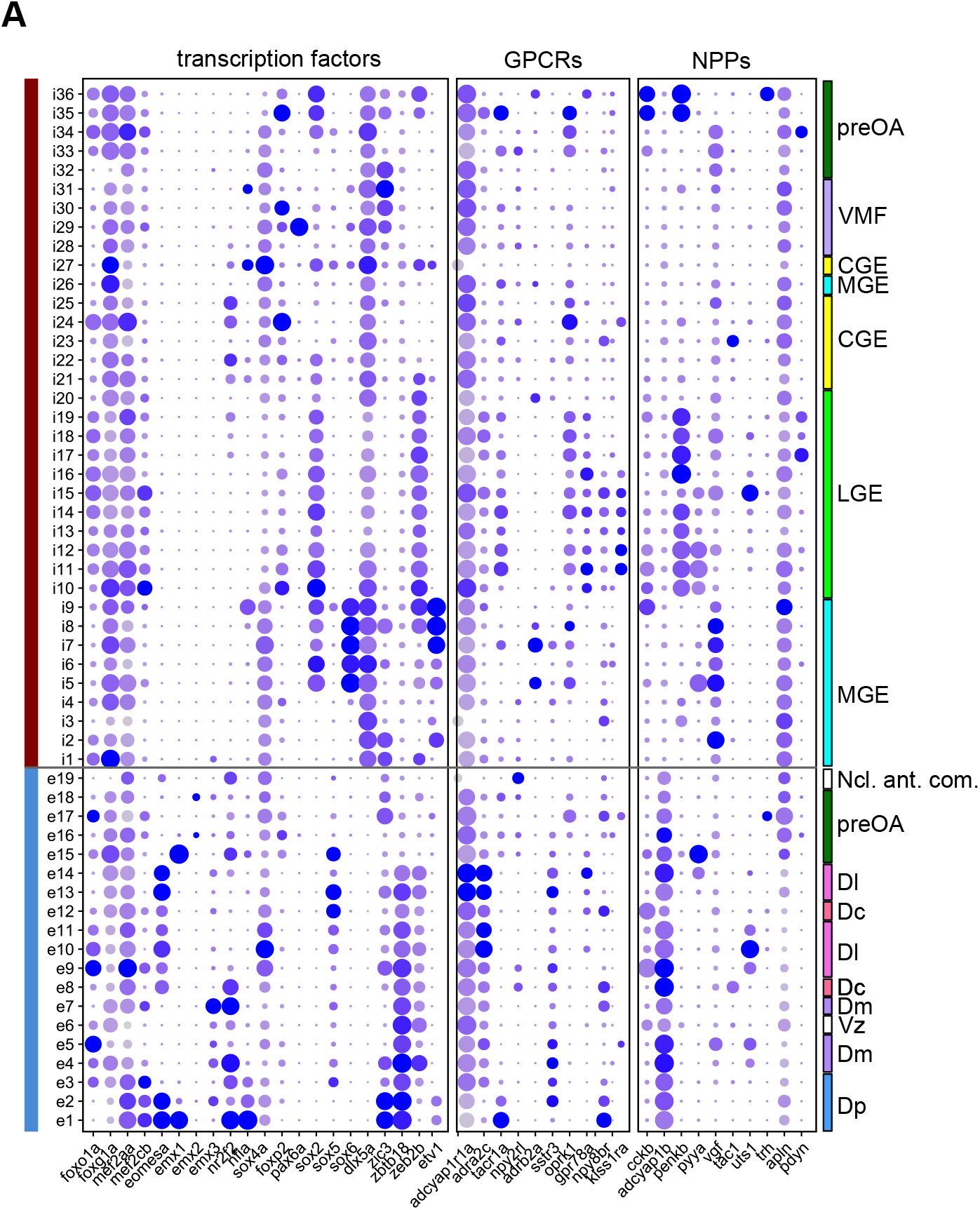
Expression pattern of marker genes. (**A**) Expression pattern of selected transcription factors, GPCRs and neuropeptide precursor genes (NPPs). Color indicates expression level (normalized log read count), and dot size represents fraction of expressing cells per cluster as in Fig. 2B. GPCRs, G protein-coupled receptors; NPPs, neuropeptide precursor; LGE, lateral ganglionic eminence; VMF, ventromedial forebrain; CGE, caudal ganglionic eminence; MGE, medial ganglionic eminence; preOA, preoptic area; Dp, posterior zone of the dorsal pallium; Dc, central zone of the dorsal pallium; Dm, medial zone of the dorsal pallium; Dl, lateral zone of the dorsal pallium, Ncl. ant. com., nucleus of the anterior commissure; Vz, ventricular zone.

As observed for TFs, high diversity of gene expression across cell types was also found for GPCRs (Fig. 3A, S3A, D, G), consistent with results from other species^27^. A cell type classifier based on GPCR expression was slightly less accurate than a TF-based classifier but still recovered the putative cell type structure of the dataset (Fig. S4D). Ribosomal genes, in contrast, were broadly expressed and did not contain sufficient information to classify cell types (Fig. S4E, H). Hence, combinatorial expression of both TFs and GPCRs provide molecular fingerprints to identify neuronal cell types.

### Localization of neuronal cell types in the pallium: putative excitatory neurons

To fully leverage our gene expression datasets, we localized cell types within the forebrain by combining bioinformatic and anatomical approaches. We first intersected the 2000 most variable genes in our single-cell dataset with the 5,203 genes that were differentially expressed between regions identified in the bulk dataset, resulting in an overlap of 734 genes. We then assumed that the gene expression profile of neuroanatomical regions is a linear combination of cell type-specific transcriptional programs. As a consequence, gene expression profiles of cell types should be correlated to gene expression profiles of brain regions where these cell types are abundant. We computed the correlation between the expression profile of individual cell types and brain regions based on the 734 genes in the intersection of datasets (Fig. 4A). Additionally, we used multi subject single-cell deconvolution (MuSIC), an approach that utilizes cell-type specific gene expression to estimate cell type proportions from bulk RNA-seq data within individual tissues. The results of this method supported those from the correlation-based analysis^43^ (Fig. S4A). These approaches are naturally limited by the precision of dissection but can serve as a first step towards cell type localization.

**Figure 4.**
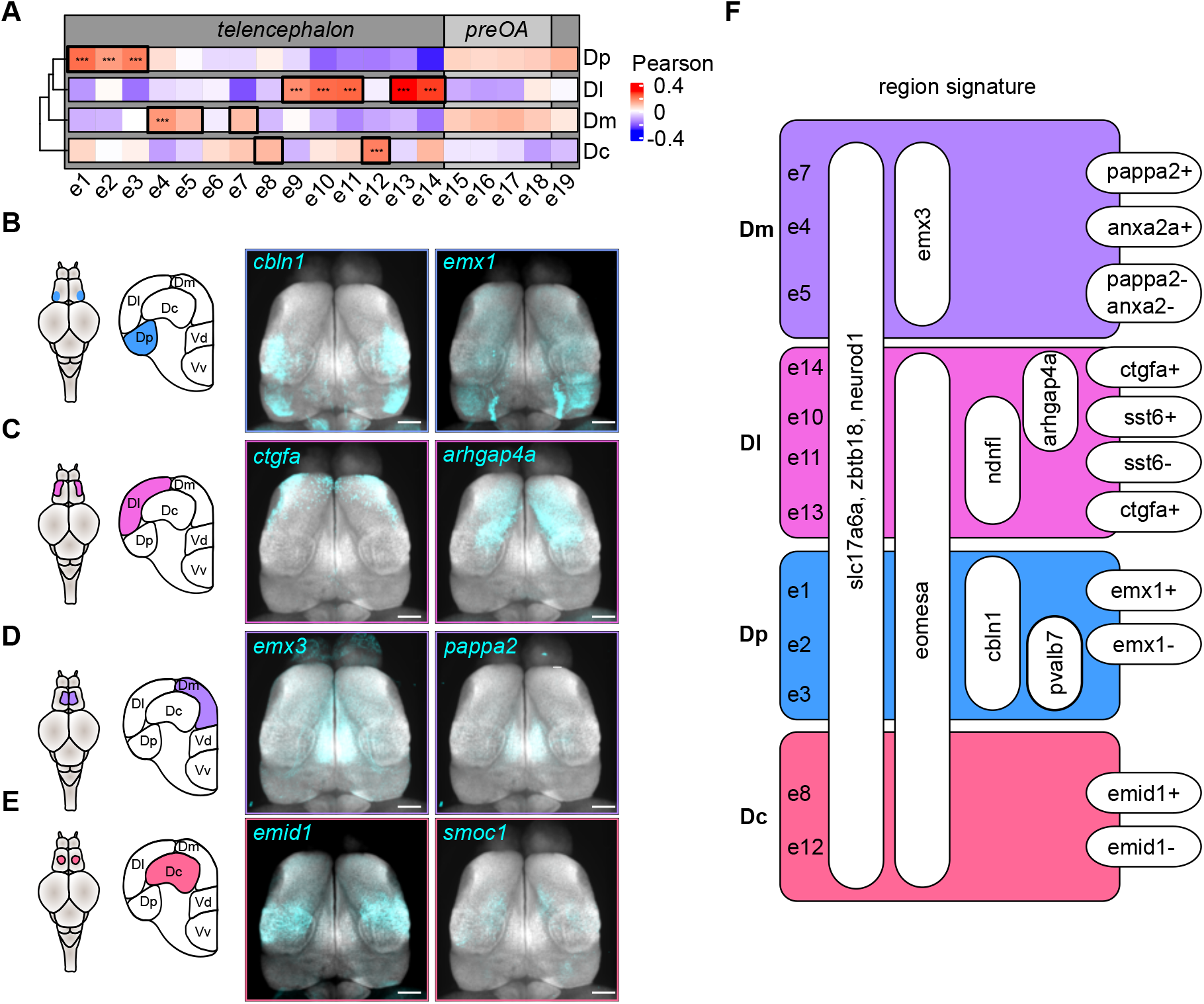
Glutamatergic cell types display anatomically discrete localization. (**A**) Correlation between the gene expression profile of individual cell types and the bulk transcriptome of pallial regions. The analysis included 734 genes that were differentially expressed between brain regions in the bulk sequencing dataset and among the 2,000 most variable genes of the scRNA-seq dataset. (**B-E**) Maximum intensity projection of gene expression patterns along the dorso-ventral axis obtained from hybridization chain reaction (HCR) against two exemplary marker genes per brain region (cyan; z-projection of confocal images) overlayed on the *snap25a* registration channel (gray). Left: location of the four pallial brain areas in a dorsal view and coronal sections of the adult zebrafish brain. Scale bar: approximately 200 pm (not precise due to registration of image datasets onto the reference brain). (**F**) Proposed hierarchical molecular signature distinguishing glutamatergic cell types in the dorsal pallium.

In a second step, we then performed whole-brain hybridization-chain reaction (HCR) in the adult telencephalon to precisely define the anatomical locations of these cell types based on marker expression^44^. This anatomical information allowed us to verify the selectivity of regional markers and to identify specific features of spatial expression patterns.

Most glutamatergic cells were found in the pallium, which is characterized by the expression of genes such as *emx1*, *emx2*, *emx3*, and *eomesa*^26^, whereas the bulk of GABAergic cells was localized in subpallial regions expressing *dlx2a* and *dlx5a*^45^.

We first focused on putative excitatory cell types and found three distinct cell types (e1-3) that were associated with Dp, the homologue of piriform cortex. These cell types could be delineated by the combinatorial expression of *emx1*, *cbln1*, and *pvalb7* (e1: *emx1*^+^/*cbln1*^+^/*pvalb7*^-^, e2: *emx1*^-^ /*cbln1*^+^/*pvalb7*^+^, e3: *emx1*^-^/*cbln1*^-^/*pvalb7*^+^, Fig. 4B, F). *Emx1* is a known marker for Dp in fish^26^ and the mammalian *emx1* lineage forms ectopic piriform cortex in *lhx2* deficient animals^46^. Expression of *cbln1* matched the subregion defined as Dp in the zebrafish brain atlas^12^, and the *emx1* expression domain demarcated the posterior part of the *cbln1* domain (Fig. 4B). These areas coincide with functionally defined subregions of Dp referred to as posterior Dp (pDp, e1: *emx1*^+^/*cbln1*^+^), a recurrent network with balanced excitation and inhibition that strongly responds to odors, and anterior Dp (aDp, e2: *emx1*^-^/*cbln1*^+^), an odor-responsive subregion driven primarily by external input^16^. The expression domain of *pvalb7* has previously been localized to the posterolateral edge of the dorsal pallium, adjacent to the region described as Dp in the atlas. This area has been proposed to be part of an extended olfactory integration complex^37,47^.

Dl was associated with multiple cell types that formed a coherent cluster in UMAP space (Fig. 2C; 4A, C, F). All cell types in this cluster (e10, e11, e13, e14; Fig. 2C) expressed *eomesa*, consistent with previous observations in Dl^26^. The region referred to as Dl in the zebrafish brain atlas can be further subdivided based on the expression of *ndnfl* (in the medial part of Dl, Fig. S7), *arhgap4a* (in posterior Dl; Fig. 4C), and *ctgfa* (in an anterior-lateral domain; Fig. 4C). Further cell types are distinguished by *sst6* (Fig. 4F).

Previous reports indicate that Dm is characterized by the expression of *emx3* and the absence of *eomesa*^26,48^, which is abundantly expressed in other pallial areas. Our correlation-based approach identified one glutamatergic cell type (e4) that was strongly associated with Dm (Fig. 4A) and embedded within an *eomesa^-^*/*emx3^+^* cluster (e4-7; Fig. 2C). In addition, two additional cell types (e5, e7) from the same *eomesa^-^*/*emx3^+^* cluster were more weakly associated with Dm (Fig. 4A). These three cell types could be further distinguished based on the expression of *pappa2* and *anxa2a* (Fig. 4D, F). The fourth cell type of the *eomesa^-^*/*emx3^+^* cluster (e6) expressed *atp1a1b*, suggesting that it contains ventricle-lining newborn neurons adjacent to Dm^49^. Spatially, we found the *emx3* domain demarcating Dm to extend into a posterior part that expresses *pappa2* while the most lateral part was characterized by expression of *anxa2a*.

Dc has been established as a separate histogenetic unit within the pallium and was proposed to be related to mammalian isocortex^13^. Previous work proposed the genes *emx2* and *eomesa* as markers for Dc^26^. Our correlation-based approach identified two cell clusters as potentially contributing to this area (e8, e12; Fig. 4E, F). While these clusters expressed *eomesa*, we did not detect *emx2* at significant levels, consistent with another single-cell sequencing study that failed to detect cell types co-expressing *eomesa* and *emx2*^37^. While cell types associated with Dp, Dl or Dm were organized in distinct neighborhoods in UMAP space, clusters associated with Dc were either adjacent to Dm-related clusters or interspersed with the clusters representing Dl (Fig. 2C). Moreover, positive correlations were observed between Dc and additional cell types, although these correlations were not significant. These observations suggest that Dc is potentially more heterogeneous in its cell type composition than other areas. Cell type e8, which was selectively correlated with Dc, was characterized by *emid1*, although this gene was also found in other cell types. The *emid1* expression domain extended from the mediolateral part of the pallium to the lateral edge of Dm, encompassing the area usually designated as Dc but also adjacent areas. The expression of the marker gene *smoc1* was more consistent with the common anatomical definition of Dc. The remaining glutamatergic cell types were part of the preoptic area, as characterized by the expression of *otpa* and *otpb*^50^, and not analyzed further.

A selection of molecular markers for the major subregions of the dorsal pallium (Dm, Dl, Dp, Dc) and their subdivisions are summarized in Fig. 3F. Moreover, Fig. S7 shows expression patterns and UMAP mappings of 43 genes that are differentially expressed in putative inhibitory and excitatory neuronal cell types in the telencephalon. These data may serve as a resource for further classifications of cell types and brain areas in the zebrafish pallium.

### Interneurons in the telencephalon

GABAergic neurons are located primarily in the ventral telencephalon (subpallium)^51^. To classify these neuron types, we first associated putative GABAergic neurons with their domain of origin based on transcription factor expression. Cells derived from the medial ganglionic eminence (MGE, i1-9, i26) were organized in three main clusters in UMAP space, which were jointly characterized by the expression of *lhx6* (Fig. 5A, E). One of these clusters co-expressed *gbx1* and *lhx8a* (i1-4, i26), suggesting that this cluster contains cholinergic neurons^9,37^. Visualization by HCR localized these cells to the medial subpallium (Fig. 5A). The second cluster contained somatostatin-positive interneurons (*sox6*+, i5-8) that were further subdivided into cell types expressing exclusively *sst1.1* or additionally *sst1.2*. The third cluster was clearly distinct from all other GABAergic types and expressed *wnt16*, *sox6, zic1*, and *nxph1* (i9; Fig. 5E, F, G).

**Figure 5.**
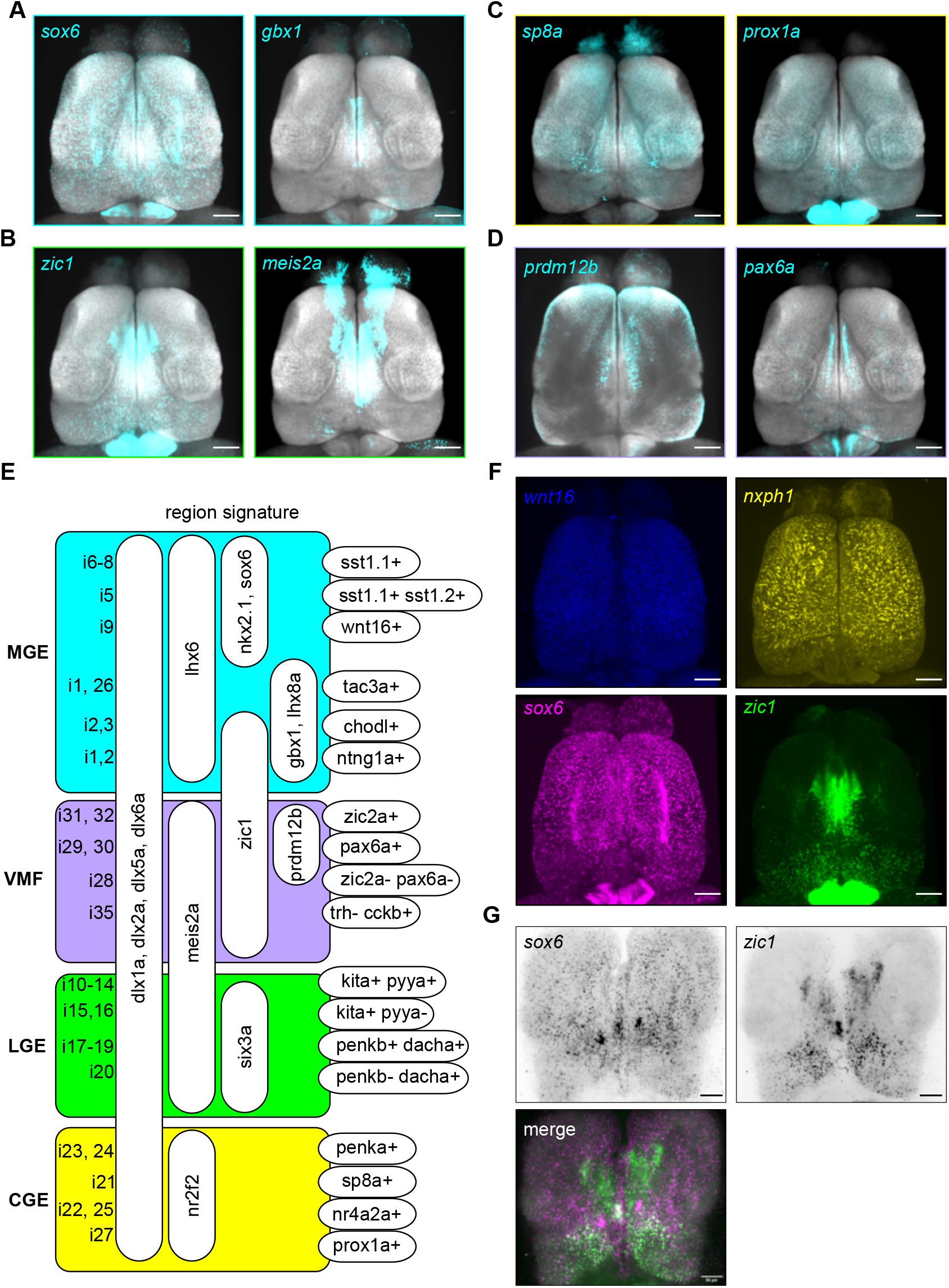
GABAergic cell types diversity in pallium and subpallium. (**A**) HCR visualization of *sox6* and *gbxl*, two marker genes for discrete medial ganglionic eminence (MGE)-derived GABAergic cell types (dorsal view, maximum intensity projections in all panels). (**B**) Cells derived from the lateral ganglionic eminence (LGE) are marked by the presence of *meis2a* and the absence of *zicl*, forming a cellular stream from subpallium to the olfactory bulb. (**C**) Exemplary marker genes of the caudal ganglionic eminence (CGE), *sp8a* and *proxla*, were expressed in a sparse band in the posterior subpallium. Outside the telencephalon, strong expression of *proxla* is seen in the habenula. (**D**) Images show *prdm12b*_+_ and subsets of *pax6a*_+_ interneurons in the ventromedial forebrain lateral to the MGE-derived *gbxl*_+_ population. (**E**) Hierarchical molecular signature of putative GABAergic cell types. (**F**) Expression of markers for MGE-derived pallial interneuron type i9 (*wnt16*, *nxphl*, *sox6*, *zicl*) is sparse and differs along the anterior-posterior axis. (**G**) Expression of *sox6* and *zicl* in different neurons, demonstrated by co­labeling in the same brain, indicating that pallial interneuron type i9 comprises multiple subtypes. Scale bars: approximately 200 pm (A-D, F); 50 pm (G).

Markers for i9 were expressed in somata dispersed throughout the pallium, as shown by HCR (Fig. 5F), indicating that interneurons from the MGE integrate into circuits in multiple pallial areas. Different marker genes showed somewhat different expression patterns: while *wnt16* was evenly distributed throughout the pallium, *nxph1* was predominantly found anterior-laterally. *Sox6* mirrored *wnt16* expression in the pallium except for a posterior region that was also occupied by *zic1* (Fig. 5F). Visualizing *sox6* and *zic1* in the same brains identified only a few cells that co-expressed both genes (Fig. 5G). In UMAP space, discrete subregions within the i9 cell cluster exhibited different gene expression patterns. The i9 cluster therefore represents pallial interneurons that are likely to comprise multiple subtypes. However, we did not attempt more fine-grained clustering because the total number of cells in this cluster was low (n = 150).

Cell types derived from the lateral ganglionic eminence (LGE, *meis2a*+, *zic1*-, i10-20) are known to migrate along a ventral stream to the striatum and olfactory bulb (OB)^52,53^. Consistent with this migratory stream, we found a band of continuous *meis2a* expression between the ventral subpallium (Vv) and the OB, supporting the notion that the migration pattern as well as the brain areas receiving interneurons from the LGE are conserved across vertebrates (Fig. 5B, E).

In addition, we found cell types that could not be associated with the LGE or the MGE but expressed genes typically found in the caudal ganglionic eminence (CGE, abundantly *nr2f2*, in subclusters *sp8a* and *prox1a*, Fig. 5C, i21-27). Markers for these cell types were found in the posterior subpallium, suggesting that all ganglionic eminences contribute to specific and distinct subpallial or pallial areas. Furthermore, some cell types could be linked to the ventromedial forebrain (VMF, Fig.5D, i28-32, i35) based on the expression of *zic1*, which was expressed in the medial subpallium, or to the preoptic area based on the expression of *galn* or *trh* (i33, i34, i36). Markers for the hierarchical classification of GABAergic cell types are summarized in Fig. 5E and additional expression patterns are shown in detail in Fig. S7.

### Evolutionary conservation of telencephalic cell types

Phylogenetic relationships between neuronal cell types across species can provide insights into conserved design principles and functions of neuronal circuits. Transcriptomics datasets from different species and developmental stages provide the opportunity to identify conserved cell types of the telencephalon by comparative genomic analyses^38–40^. As gene-regulatory programs diversify over evolutionary timescales, the identification of homologous cell types is expected to become more difficult with increasing phylogenetic distance. In zebrafish, an additional challenge arises from the recent duplication of the genome^41,54^, which often results in an accelerated evolution of one paralog^49^. Previous approaches to identify conserved cell types typically relied on one-to-one orthologous genes and co-clustered cell types based on conserved correlation structure^40,55^. This strategy has been successful in amphibians, reptiles, and mammals^27,38^ but presents challenges in zebrafish because genome duplication substantially reduces the number of clearly defined one-to-one orthologs. Previous studies thus compared zebrafish to rodents or human by either adding artificial genes to account for the large number of one-to-many orthologous^28^ genes or by averaging the expression across duplicated genes^37^. Here we applied a method that maps gene expression datasets from different species onto on a self-assembling latent manifold^54^. The approach includes an initial manifold alignment based on sequence similarity of homologous genes and further alignment steps based on gene-gene correlations until convergence. This method has been designed specifically to identify related cell types across species and does not require the definition of arbitrary rules for one-to-many mapping of orthologous genes. We therefore expected this method to detect phylogenetically conserved cell types with higher specificity than other approaches. We compared the cell types identified in our study to a dataset from the entire brain of a lizard, the bearded dragon (*Pogona vitticeps*)^27^. This dataset was chosen because it has already been analyzed extensively regarding cell type conservation, thus allowing for further conclusions across species. Moreover, as this dataset encompassed the entire CNS, it offered opportunities for internal controls by comparing telencephalic cell types of zebrafish to cells in the telencephalon and other brain regions of the lizard.

Distinct cell types such as ependymoglial cells or microglia could be mapped specifically and reliably between zebrafish and the bearded dragon (Fig. S5A). We therefore proceeded to identify conserved neuronal subtypes. As expected, GABAergic and glutamatergic cell types in the zebrafish telencephalon achieved highest alignment scores with neurons from the lizard telencephalon (Fig. 6A). Outside the telencephalon, the best match was found for habenular cells, which we included in this analysis as the habenula is well-conserved across species^27^. Among the glutamatergic cells in the zebrafish forebrain, Dl-associated cell types (e10, e11, e13, e14) showed high alignment scores specifically to cell types from the medial (MC) and dorsomedial lizard cortex (DMC) (Fig. 6B). Topological analysis of the zebrafish telencephalon suggested that Dl is homologous to the medial pallium in other species, which corresponds to MC/DMC in lizards and the hippocampal formation in mammals^13^. As previous work has linked cell types of MC to the mammalian dentate gyrus and cells from DMC to CA1 and CA3^40^, we investigated whether cell types within Dl are spatially segregated. We found that cell types associated with DMC exclusively expressed the gene *ctgfa*. The expression of this gene was restricted to anterior Dl, suggesting that this area may be homologous to DMC and, by extension, to mammalian CA1 and CA3 (Fig. 6C) at the cell type level. When we determined the transcription factors driving this cross-species alignment, we identified *neurod1* and *neurod2* as a common core set of transcription factors in the entirety of Dl (Fig. 6D). These genes are enriched in rodent hippocampus, cerebellum, and cortex^56^.

**Figure 6.**
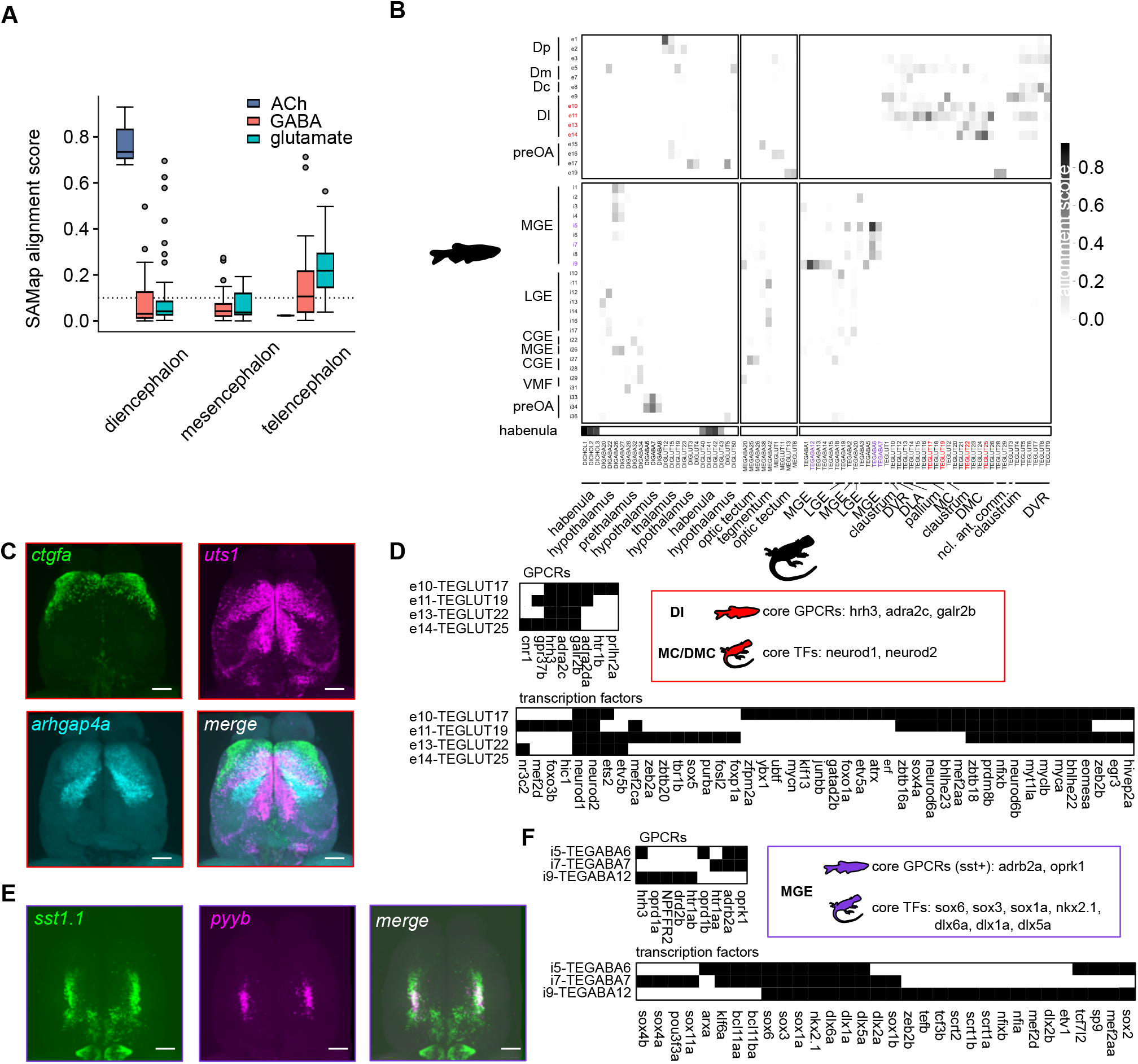
Evolutionary conservation of cell types between fish and reptiles. (**A**) Scores of cell type alignments between the zebrafish telencephalon and the bearded dragon whole brain atlas, subsampled for telencephalic, mesencephalic and diencephalic neuronal cell types, segregated by their neurotransmitter (ACh was not detected in the mesencephalon). The analysis was restricted to zebrafish cell types with a minimum alignment score of 0.1 (dotted line). ACh, Acetylcholine. (**B**) Alignment scores of individual cell types. Blocks separate glutamatergic and GABAergic cells in the zebrafish (vertical) and brain regions in the lizard (horizontal). Color-coded cell types are used as examples for alignment analysis in (C-F). (**C**) Expression of markers for cell types in the zebrafish telencephalon that were associated with MC and DMC in the lizard (dorsal view, maximum intensity projections; same in all panels). Expression patterns stratify Dl along the anterior/lateral-to-posterior/medial axis. The marker gene *ctgfa* is expressed in an anterolateral domain that was associated by cell type alignment with DMC in lizards, which has been linked previously to hippocampal regions CA1 and 3 in mammals. Posteromedial regions expressing *utsl* or *arhgap4a* were associated with MC in lizard, which has been linked to the mammalian dentate gyrus. (**D**) Binarized gene expression patterns of conserved GPCR and transcription factors in Dl of zebrafish and MC/DMC of the lizard. (**E**) Overlapping of *pyyb* and *sstl.1* expression in the zebrafish telencephalon, demonstrating that *pyyb* represents a subtype of *sst*_+_ cell types, which are conserved in the lizard. (**F**) Binary expression patterns showing a common transcription factor code for MGE-derived neurons in zebrafish and the lizard, but distinct GPCR fingerprints of *sst*_+_ and *wnt16*_+_ interneurons. Scale bars approximately 200 pm.

Dc in zebrafish has been suggested to be homologous to isocortex. In our analysis, a cell type associated with Dc (e8) aligned well to the glutamatergic neuron type TEGLUT8 in the dorsoventricular ridge (DVR) of the lizard, which is a multimodal sensory-recipient telencephalic structure in birds and reptiles (Fig. 6B)^57^. Although TEGLUT8 is transcriptomically similar to excitatory pyramidal neurons in layer 4 of the mammalian neocortex^27^, it has been suggested that these cell types arose by convergent adaptation of gene modules associated with sensory information processing^40^.

Among the GABAergic cell types in the zebrafish telencephalon, only MGE-derived cell types showed considerable alignment with neurons in the bearded dragon. In particular, somatostatin-positive interneurons (i5-8) aligned with the sst-expressing MGE-derived neurons in the lizard telencephalon (Fig. 6B, E). In zebrafish, the somatostatin1 gene has two paralogs that are expressed in separate but partially overlapping subpopulations of the subpallium (sst1.1: i5-8, sst1.2: i5; Fig. 6E). While all of these neuron types were most closely aligned to the lizard cell type TEGABA6 (SST-RELN), the transcriptome of i5 additionally aligned with other MGE-derived sst+ cell types (Fig. 6B), suggesting that the sst1.2+ population has adapted additional genetic modules. The GABAergic interneuron population in the pallium (i9; Fig. 5F, G) aligned strongly with multiple distinct cortical interneuron types (COL19A1, LAMP5, CRHBP, SST-CORT), further indicating that this small cluster comprises multiple cell types. The set of transcription factors driving the cross-species alignment for MGE-derived cell types contained the well-conserved marker genes *sox3*, *sox6*, and *nkx2.1* (Fig. 5E, 6F). We further observed that galanin-positive GABAergic hypothalamic cells of the lizard aligned with galanin-expressing neurons from the zebrafish preoptic area (i33, 34, Fig. 6B, S5B).

To analyze whether combinatorial expression of GPCRs was retained through evolution we extracted the set of genes that were differentially expressed and significantly increased the cross-species correlation between cell types. This analysis revealed that the expression of a subset of GPCRs was retained between related cell types in zebrafish and the bearded dragon (Fig. 6D, F, S5B). For example, expression of the galanin receptor *galr2b*, the adrenergic receptor *adra2c*, and the histamine receptor *hrh3* was detected in all cell types of Dl. Expression of these genes contributed significantly to the alignment with MC or DMC-related cell types in the lizard, indicating that orthologous genes remained co-expressed in this structure over evolutionary timescales and that evolutionary pressure kept these modulatory systems in place, suggesting conservation of information processing within the amniote medial cortex.

In summary, some cell types are highly conserved between zebrafish and lizards while others cannot be mapped to putative sister cell types. Similar general observations were made in comparisons between other vertebrate species^27,38,39^. The cell types that we found to be conserved between fish and reptiles are also conserved between reptiles and mammals^40^, suggesting that there is a core set of common neuronal cell types in vertebrates. In addition, a significant set of neuron types may be derived from common ancestors but are not identified as homologous because they developed species-specific novelties or underwent steady transcriptional adaptation.

### Cell type-specificity of neuromodulatory signaling components

The identification and localization of cell types in the telencephalon allowed us to directly examine the combinatorial expression of markers for different neuromodulators and their receptors across neurons and brain regions. To analyze potential receivers of neuromodulatory signals we first binarized the expression of all metabotropic receptors for the four “classical” neuromodulatory systems (cholinergic, dopaminergic, norepinephrinergic, and serotonergic GPCRs) and determined the number of cell types expressing each of these receptors. All neuronal cell types in our dataset expressed receptors of at least three systems (Fig. 7A, B), implying that most or all neuron types in the zebrafish telencephalon are susceptible to modulation by multiple classical neuromodulators. Different cell types expressed unique combinations of GPCRs, indicating that combinatorial actions of neuromodulators may differ between cell types (Fig. 7A). On average, GABAergic cells expressed more GPCRs than glutamatergic cells (classical modulatory systems in Fig. 7B, neuropeptidergic systems in Fig. S6A), indicating that the modulation of inhibitory interactions plays an important role in the control of neural circuit function.

**Figure 7.**
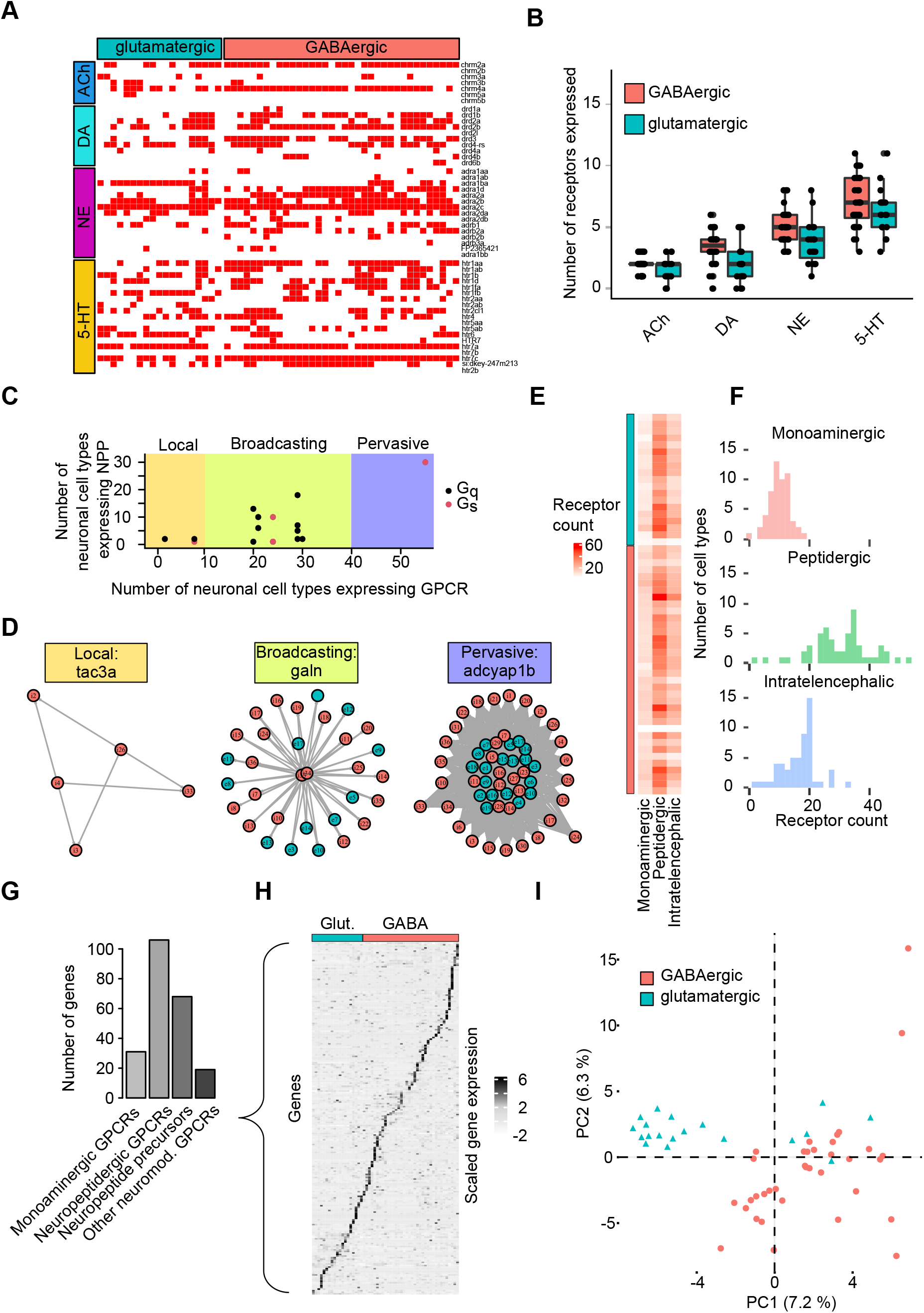
Neuromodulatory networks within the zebrafish telencephalon. (**A**) Binarized expression of monoaminergic GPCRs (y-axis) in different cell types (x-axis). Expression was counted if reads were detected in at least 5% of cells per cell type. (**B**) Number of GPCRs for different monoaminergic neuromodulators expressed by glutamatergic and GABAergic cells. Boxplots show median, hinges correspond to first and third quartile. Whiskers extend to the largest value no further than 1.5 * the inter-quartile range. Dots show individual data points. (**C**) Relationship between number of cell types expressing NPPs and their receptors. Each data point represents one NPP-receptor pair. Shadings indicate regions corresponding to different types of networks (“Local”, “Broadcasting”, “Pervasive”). Most networks are of the “broadcasting” type. G protein type indicated by color. (**D)** Graphs show individual examples of each type of signaling network. (**E**) Number of GPCRs for three signaling systems in all telencephalic cell types. (**F**) Distributions of receptor counts shown in (E). (**G**) Number of expressed genes associated with neuromodulation, separated into four different categories. (**H**) Normalized (z-scored) expression levels of all expressed genes associated with neuromodulation, sorted by maximum expression per cell type. See Fig. S7 for an enlarged display. (**I**) Normalized expression matrix in (H) projected onto the first two PCs. GABAergic and glutamatergic cell types are separated but no obvious additional clusters are observed.

We next extended our analysis to some of the most abundantly expressed neuropeptides and their receptors. As observed in telencephalic neurons of rodents and other vertebrates^6,22,58^, we found that all cell types express neuropeptide precursors (NPPs), with co-expression of multiple NPPs being the rule rather than the exception (Fig. S6A). Some GABAergic cell types co-expressed combinations of NPPs that were also co-expressed in homologous cell types of other species. For example, the co-expression of somatostatin and neuropeptide Y (*npy*) in a subset of MGE-derived sst^+^ cells was observed in the zebrafish telencephalon (Fig. S6A), the turtle pallium, the songbird DVR, and the mouse and human motor cortex^22^.

To better understand the directionality of information flow we binarized expression of components of all neuropeptidergic systems and selected those for which we could identify both ligand and receptor-expressing neuron classes in our dataset (Fig. 7C). Based on the resulting connectivity matrices, we visualized relationships between ligand- and receptor-expressing neuron classes in network graphs (Fig. 7D) and classified network topologies into three groups^59^: (1) “Local” networks, in which both the ligand and the receptor were only expressed in a few cell types (Fig. 7C, D); (2) “Broadcasting” networks, which contain a small number of “sender” cell types and a large number of potential “receiver” (receptor-expressing) cell types; and (3) “Pervasive” networks with a many-to-many signaling structure. Most networks were of the “broadcasting” type (Fig. 7C, D), while “local” networks were rare and only one network was “pervasive”. The latter was the signaling network of the adenylate cyclase activating polypeptide 1b (*adcyap1b*), the most widely expressed neuropeptide in the zebrafish telencephalon (Fig. 7C, D, S6A).

To estimate the number of modulatory networks that a given cell type is embedded in we calculated the number of receptor types expressed by each neuron class. We stratified the data into monoaminergic and neuropeptidergic GPCRs, which we further stratified into either all receptors or the subset for which the respective ligand was expressed in our telencephalic dataset (intratelencephalic signaling network; Fig. 7C). This analysis revealed that individual cell types expressed on average 10 monoaminergic receptors and about 30 peptidergic GPCRs (Fig. 7E, F), out of which approximately 20 were part of intratelencephalic signaling networks. These results suggest dense neuromodulatory interactions between telencephalic neuron types^21,22^.

In other species such as nematodes, neuromodulatory networks exhibit a mesoscale architecture with distinct groups of cell types that share a similar input-output structure^59^. To examine whether a similar organization exists in the zebrafish telencephalon we compiled a gene list containing all NPPs and neuromodulatory GPCRs expressed in our single-cell sequencing dataset (Fig. 7G). We then analyzed expression not as binary patterns but quantified the strength of gene expression by the normalized (z-scored) number of reads across all cell types. Although expression of individual genes was usually detected in multiple cell types, high expression was typically detected only in one or a few cell types (Fig. 7H). Consequently, each cell type could be distinguished by the specific high-level expression of one or a few neuromodulatory genes. This conclusion was confirmed by the observation that individual principal components of the normalized (z-scored) gene expression matrix explained only relatively small fractions of the total variance (PC1: 7.2%; PC2: 6.3%, Fig. 7I). Projecting cell types onto the first two PCs of the normalized gene expression matrix separated glutamatergic and GABAergic cells (Fig. 7I), consistent with systematic differences in gene expression between major classes of neuronal cell types. However, no obvious further clustering of cell types was found in this space, supporting the notion that different neuron types are likely to exhibit unique patterns of neuromodulatory interactions. Neuronal cell types are therefore embedded in many (typically tens of) modulatory networks (Fig. 7A-H), but their expression profile may favor signaling through a small subset of modulators (Fig. 7H, I, S6B, C).

## Discussion

We used transcriptomics and HCR to delineate cell types and neuromodulatory networks in the telencephalon of adult zebrafish. Bulk sequencing of micro-dissected tissue revealed distinct molecular signatures of pallial subregions including specific patterns of GPCR expression. Single-cell sequencing at high depth identified 55 neuronal cell types and associated them with distinct telencephalic areas based on molecular fingerprints and marker gene visualization. The results provide new insights into the organization of the zebrafish telencephalon and into phylogenetic relationships between cell types and brain areas across species. Targeted analyses of genes associated with neuromodulation indicate that different neuromodulatory systems act in concert and provide insights into the organization of the underlying networks. Moreover, our data provide resources for the manipulation of specific cell types and brain areas and for analyses of phylogenetic relationships across species.

### Brain areas and cell types

The molecular characterization of cell types and the spatial mapping of many marker genes largely confirmed the boundaries between major subregions of the dorsal telencephalon (Dp, Dl, Dm, Dc) and revealed additional anatomical substructures (Fig. S6)^14,26^. For example, within Dp, we found a segregation of cell types (e1: *emx1*^+^/*cbln1*^+^; e2: *emx1*^-^/*cbln1*^+^) between posterior and anterior compartments that coincide with the subdivision into pDp and aDp, respectively, based on electrophysiological observations^16^. These observations support the hypothesis that pDp and aDp perform different computational functions. Moreover, spatially distinct expression patterns of additional Dp-related marker genes (e.g., *lhx9*, *nfia*) indicate additional functional subcompartments within pDp.

Dl has previously been divided into dorsal (Dld) and ventral (Dlv) parts based on embryonic development^51^ and behavioral observations in other teleosts^60^. However, we observed no prominent dorso-ventral asymmetries in gene expression but a segregation of Dl-related cell types along a posterior/medial-to-anterior/lateral axis (posteromedial: e10: *ndnfl^+^/ arhgap4a^+^/ sst6^+^/ ctgfa^-^*; e11: *ndnfl^+^/ arhgap4a^-^/ sst6^-^/ ctgfa^-^*; anterolateral: e13: *ndnfl^+^/ arhgap4a^-^/ sst6^-^/ ctgfa^+^*; e14: *ndnfl^-^/ arhgap4a^+^/ sst6^-^/ ctgfa^+^*;). To our knowledge, it is currently unknown whether this pattern reflects a functional segregation of neuronal circuits within Dl. Well-defined subregions were also identified within Dm based on the expression of markers for Dm-related cell types (e.g., *pappa2* in the posterior part of Dm; see Fig. S6 for details). Dc was the only area in the dorsal pallium where distinct cell types and well-defined subregions were not clearly delineated, possibly because the resolution of cell types is complicated by high heterogeneity and low abundance.

Phylogenetic relationships between cell types were explored using an approach that maps transcriptomic fingerprints onto a self-assembling manifold (SAMap)^54^. Unlike classical approaches, this algorithm uses the full transcriptome of zebrafish cells despite the recent genome duplication^54^. Strong alignments were found between a subset of neuronal cell types in zebrafish and a reptile, the bearded dragon^27^ and to other vertebrate species by extension. These observations suggest that vertebrates contain a conserved set of common cell types while other cell types evolved more idiosyncratically. Molecular fingerprints of conserved cell types can therefore help uncover phylogenetic relationships between brains that develop by evagination or eversion^13,41,54^.

Our results support the previously established link between Dl and hippocampal areas, a relation between Dc and isocortex, and the conservation of sst+ interneurons. In addition, alignments of cell types associate Dl with areas MC and DMC in the lizard, which have been linked to the dentate gyrus and hippocampal areas CA3 and CA1, respectively^40^. Consistent with this observation, lesions of Dl (primarily Dlv) impaired map-based spatial navigation and trace conditioning in goldfish, consistent with effects of hippocampal lesions in mammals^60–62^. More detailed cell type mapping suggests that neuronal circuits related to DM (dentate gyrus) are located in posterior-medial Dl (cell types e10, e11) while circuits related to DMC (CA1/3) are located in anterior-lateral Dl (e13, e14). Recordings in behaving goldfish revealed neurons with functional properties related to navigation (e.g., putative speed cells, border cells, head direction cells)^63,64^ but equivalents of grid cells or place cells have not been described. The molecular characterization of subregions in Dl may thus inform studies of cognitive maps in teleosts.

Among GABAergic cell types, strong alignment was observed between interneurons derived from the MGE, which localize primarily to the subpallium. However, one MGE-derived interneuron type (i9) was found at low density throughout the dorsal pallium. Markers such as *nxph1* and *zic1* revealed differences between individual i9 cells and systematic changes along the anterior-posterior axis, suggesting that this interneuron class can be further subdivided. Our results may thus provide a starting point for a more detailed intra- and inter-species analysis of pallial interneurons^66,67^, which are increasingly moving into the focus of physiological studies.

### Neuromodulatory networks in the zebrafish telencephalon

A first step towards the systematic characterization of neuromodulatory networks is to determine the number and combination of neuromodulators that can act upon different cell types. We found that telencephalic cell types in zebrafish typically express diverse and unique sets of neuromodulatory GPCRs, and that most cell types are embedded in approximately 30 - 40 monoaminergic and peptidergic signaling systems. As a corollary, the combinatorial expression of neuromodulatory GPCRs can serve as a molecular fingerprint for individual cell types. Similar general observations were made in nematodes^3,59,68,69^ and other vertebrate phyla^21,22,58^. These results support the notion that neuromodulators typically act in combination to control neuronal circuit function. However, because high-level expression of neuromodulatory genes across cell types is sparse, modulatory effects on individual cell types may be dominated by one or a few systems. Furthermore, our results indicate that many neuromodulatory systems exhibit a broadcasting organization where a small number of senders can potentially influence a large number of receivers. This architecture is generally consistent with global functions of neuromodulation such as the control of brain states.

The systematic analysis of expression patterns at single-cell resolution provides a basis to decipher the logic of combinatorial neuromodulation by identifying cell types that act as potential “senders” and “receivers” of neuromodulatory signals. This information may now be complemented by activity measurements and functional manipulations to explore how multiple neuromodulatory signals are integrated to control internal brain states and behaviors^7,70^. While synergistic interactions between neuromodulators have been described^18,21,73,74^, a comprehensive understanding of such interactions remains elusive. Our phylogenetic analysis revealed that homologous cell types retained a subset of co-expressed GPCRs over evolutionary time scales, such as the galanin receptor *galr2b* and the histamine receptor *hrh3* in cell types associated with Dl. While galanin dampens hippocampal excitability in rodents^71^, histamine has the opposite effect^72^. The conserved co-expression of signaling systems for histamine and galanin in the medial cortex (Dl in fish, hippocampus in mammals) may therefore indicate that these neuromodulators interact and mediate fundamental functions in the control of brain states or memory formation^73^.

Neuropeptides and other modulators are released into the interstitial space and reach target cells via volume transmission^3^. For this and additional reasons, neuromodulation is often considered complementary to the information flow in synaptically connected networks and assumed to mediate signaling between broad classes of cells. Hence, the reconstruction of neuromodulatory networks may focus on cell types, rather than individual neurons, as functional units, and connectivity may be inferred from the expression of markers for neuromodulators and their receptors along with coarse anatomical information^3,59^. Even as adults, zebrafish provide the opportunity to optically measure and manipulate neuronal population activity in the telencephalon during behavior^8^. Unlike larvae, adult zebrafish show pronounced cognitive behaviors while the brain is still sufficiently small to enable the ultrastructural reconstruction of pallial brain areas at synaptic resolution by volume electron microscopy and automated segmentation^74,75^. The molecular analysis of neuromodulatory networks in adult zebrafish may thus be combined with other approaches to analyze mechanisms of neuromodulation that are not accessible in larger vertebrates.

## Materials and Methods

### Animal Husbandry

Zebrafish (Danio rerio) were raised and kept in 3.5 L tanks under standard laboratory conditions (26 - 27 C°; 13 h/11 h light/dark cycle, pH = 7.5 +/- 0.3, conductance = 650 μS +/-282) in groups of up to 10 animals per liter (ZebTEC Active Blue System; Tecniplast). Fish were fed age- and density-dependent amounts of GEMMA Micro (Skretting) and brine shrimp. All procedures followed institutional guidelines and were approved by the Veterinary Department of the Canton Basel-Stadt, Switzerland.

### Dissection of adult brain areas

Adult zebrafish (4 - 6 months post fertilization, mpf) were sacrificed by an overdose of tricaine methanesulfonate (MS-222) and decapitated in ice-cold (0 - 4 °C) artificial cerebrospinal fluid (ACSF, 124 mM NaCl, 2 mM KCl, 1.25 mM KH_2_PO_4_, 1.6 mM MgSO_4_, 22 mM D-(+)-Glucose, 2 mM CaCl_2_, 24 mM NaHCO_3_, pH 7.2). Brains were dissected by carefully removing the jaw, the eyes, the soft tissue surrounding the bones and the ventral bones of the skull with fine forceps. Brain used for imaging were transferred to ice-cold Formal-FIXX solution (diluted to 10% in phosphate-buffered saline with 0.1 % Triton-X 100, PBT) and kept at 4 °C on a rotating device for 48 hours. Afterwards, brains were washed 5 times with PBT and transferred stepwise to 100 % MeOH. At this stage, brains could be kept at -20°C for up to 6 months. For the microdissection of different pallial areas, we used the line Tg[vglut1:GFP;vglut2:DsRed]^76,77^ and dissected Dm, Dl, Dc, and Dp guided by the combination of gross anatomy and the joint expression of the transgenes.

### Dissociation of telencephalon

Single-cell suspensions were generated following established procedures^78^. After extracting the brain from the skull, the olfactory bulb and areas posterior to the telencephalon were removed with a scalpel. The telencephalon was transferred to Buffer Mix-I of the Neural Tissue Dissociation Kit (Miltenyi, Cat.Nr. 130-092-628) supplemented with 100 nM TTX and 4 μM Actinomycin-D. Two brains were combined per replicate. In total, four fish were used for single-cell dissociation. Brains were slowly rotated at 28.5 °C and triturated with a glass capillary to aid physical dissociation. After the addition of Buffer-Y mix, the procedure was repeated until no visible chunks were left in the liquid. A cell strainer (40 μm) was equilibrated with 5 mL of 4% BSA in PBS solution and the cell solution was passed through the strainer. 4 mL PBS were added to dilute BSA to 2%. Cells were centrifuged in a benchtop centrifuge at 300 g for 10 minutes at 4 °C. After removal of supernatant, cells were resuspended in 10 mL 2 % BSA in PBS and pelleted again. This time, cells were resuspended in 400 μL PBS and placed on ice.

### Fluorescence activated cell sorting (FACS)

To enrich living cells, we added 1 μL of DRAQ7^TM^ (ThermoFisher, Cat.Nr. D15106) dye to the cell suspension and excluded non-viable cells via flow cytometry, using a FACSAria (BD Biosciences).

### Next-generation sequencing and bioinformatic analysis

After sorting 16,000 cells per run, scRNA-seq libraries were generated using the 10X Genomics Chromium 3’ Reagents Kit (v3.1 Chemistry) and sequenced with a NovaSeq Illumina machine. Using STARsolo^79^, reads were aligned to the zebrafish genome (GRCz11) without regarding alternative loci, relying on the transcriptome annotation provided by the Lawson Lab^80^. Subsequent analysis of the scRNA-seq dataset was mainly performed using Seurat v4.3.0.1^81^. We first filtered out low-quality cells based on a combination of features (> 500 genes, < 0.1 % of mitochondrial genes per cell). Counts were normalized, scaled, dimensionality-reduced with PCA and Louvain-clustered using a resolution chosen to maximize silhouette width. This first round of clustering returned higher-order cell classes such as oligodendrocytes or GABAergic cells. We then selected neuronal cells based on the expression of the genes *snap25a*, *slc17a6a*, or *slc32a1* and subclustered GABAergic and glutamatergic cells independently. For further analysis, we compiled gene lists for different gene families (transcription factors, GPCRs, cell adhesion molecules, proteoglycans, ion channels, ribosomal proteins) from the KEGG database^82^ (search terms dre03000, dre04030, dre04515, dre00535, dre04040, dre03011) and computed the cumulative expression across cell types. Additionally, we trained random forest classifiers to predict cell type identity based on individual gene families by building 500 decision trees for each gene family (R library randomForest).

For cross-species analyses, we used SAMap^54^ to relate cell types between our dataset and *Pogona vitticeps*^27^ by following established procedures. In short, we constructed a BLAST map between the nucleotide sequences of the longest transcripts per gene of zebrafish and *P. vitticeps*. We then projected both unprocessed scRNA-seq datasets into the joint latent space of the BLAST map and relied on the algorithm to expand cell neighborhoods to determine cross-species neighbors. After calculating mapping scores for cell types, we thresholded alignment at >= 0.1 for further analysis. Ultimately, we used the *GenePairFinder()* function to identify genes that were differentially expressed and contributed positively to the cross-species correlation between cell types.

To analyze neuromodulatory networks we first applied a thresholding method to determine whether a gene was expressed in a cell type. GPCRs were counted as expressed if reads were detected in 10 % of cells within a cell type cluster. This approach was used to compute binary expression matrices and calculate the distribution of GPCRs expressed per cell type. For neuropeptidergic networks, we identified GPCR-ligand pairs and computed the corresponding cell type adjacency matrices. Using the igraph library^83^, we constructed network graphs for all intra-telencephalic peptidergic signaling systems. Ultimately, we compiled a list of all genes involved with neuromodulation and scaled the expression data across all cell types for analysis with PCA. To visualize gene expression data, we sorted the matrix based on maximum expression per cell type.

For microdissected tissue, we obtained sorted cells after dissociation and prepared libraries with the SmartSeq2 protocol for pooled cells. Sequencing was performed using a HiSeq 2500 Illumina machine. Using the same STAR index as for the scRNA-seq dataset, the raw reads were aligned using the “GeneCounts” option (‘—quantMode’) to count the number of reads per gene while mapping with STAR. Bulk sequencing data was analyzed using DESeq2^84^. We identified genes differentially expressed in different regions and annotated with the ENSEMBL gene ID, Entrez gene ID and GO term using the BioMart pipeline^85^. GPCRs were identified by matching Entrez gene ID with the KEGG database (search term dre04030). For correlation analysis, we obtained the intersection of the 2,000 most variable genes in our scRNA-seq dataset and the regionally differentially expressed genes from the bulk dataset. Gene expression for scRNA-seq was averaged over cell types and averaged over regions for the bulk dataset. Both resulting matrices were z-scored to get a standardized expression level that was then used for correlation analysis. Additionally, we used the R package MuSIC to deconvolve the contribution of individual cell types to distinct pallial regions as described^43^.

### Hybridization Chain Reaction (HCR)

For HCR, MeOH-fixed brains were treated with 2 % H_2_O_2_ in MeOH for 20 minutes at room temperature. Afterwards, specimens were stepwise rehydrated by incubating for 5 minutes in 100, 75, 50, and 25 % MeOH (in PBT). After washing specimens 5 x 5 minutes in PBT, the clearing process was started. In brief, brains were incubated at 37 °C in solution 1.1 of the Deep-Clear protocol (adapted as 10% THEED, 5% CHAPS, 5% urea, pH = 11.0)_86_. Afterwards, specimens were washed in PBT and pre-hybridized with probe hybridization buffer for 30 minutes at 37 °C_44_. Probe solution was prepared by adding 1 - 1.5 μl of 250 nmol per probe stock to 500 μl of probe hybridization buffer at 37 °C. Pre-hybridization solution was then replaced by probe solution and specimens were incubated for at least 48 hours at 37 °C. Specimens were washed 4 times with probe wash buffer at 37 °C for 15 minutes each and 4 times with 5x SSCT buffer at room temperature. Brains were now placed in 350 μl of amplification buffer for at least 30 minutes. 10 pl of 3 pM hairpin stock solution was placed at 95 °C for 90 seconds and allowed to cool to room temperature in a dark drawer. Hairpin solution was now added into 500 μl of amplification buffer and the specimens were placed in this solution for at least two days at room temperature. Excess hairpins were removed by washing with 5x SSCT. For RI-matching, specimens were placed in solution 2.2 (PBS, pH = 9.0, meglumine diatrizoate, 50% w/w) immediately before imaging.

### Microscopy

Specimens were mounted dorsally and visualized with an inverted spinning disc confocal microscope (AxioObserver7, Zeiss, Jena, Germany). For all conditions, brains were imaged with a 10x air objective (NA 0.45, Plan-APOCHROMAT), using laser wavelengths 488, 561, and 640 nm (Coherent Obis).

### Mapping to reference brain

All obtained stacks were stained against *snap25a* as reference channel. Registration to one representative brain was performed with the CMTK toolkit with standard settings^87,88^.

## Supporting information

Sequence information for oligos

## Acknowledgements

We are grateful to Jan Eglinger and Laure Plantard for help with the CMTK algorithm and image acquisition, to Charlotte Soneson for help with analysis, to Hubertus Kohler for FACS sorting, to Sebastian Smallwood and Aluri Sirisha for help with sequencing, and to the Friedrich group for scientific discussions. We also thank David Hain for discussing the *P. vitticeps* dataset with us. L.A. is supported by an EMBO postdoctoral fellowship (ALTF 379-2022). This work was also supported by Novartis Research Foundation, by fellowships from the European Union (Marie Curie) and JSPS to C.S., by the European Research Council (ERC) under the European Union’s Horizon 2020 research and innovation program (grant agreement no. 742576), and by the Swiss National Science Foundation (grant no. 310030_212236/1).

## Author Contributions

Experimental design and data interpretation: L.A. and R.F. Experiments: L.A. and C.S. Bioinformatic analysis: L.A. and H.H. Manuscript writing: L.A. and R.F. Project management and supervision: R.F.

## Data Availability

Sequencing data is available from the SRA depository linked with the BioProject PRJNA964500. Previously published sequencing data is available from the Gene Expression Omnibus datasets GSM3334110^89^, GSE106121^90^, GSE105010^36^ and the NCBI sequence read archives PRJNA591493^91^ and PRJNA812380^27^.

## Code Availability

Analysis code is available from the authors and from the public repository https://github.com/Anneser/zebrafishTelencephalon.

## List of supplements

Tabe S1: list of oligo pools used for HCRs

Table S2: list of GPCRs and their normalized bulk expression

Figure S1: Bulk sequencing of pallial microdissected areas

Figure S2: Quality assessment of scRNA-seq dataset

Figure S3: Cell-type specificity of gene expression

Figure S4: Contribution of individual cell types to pallial areas

Figure S5: Evolutionary conservation of cell superclusters between zebrafish and bearded dragon

Figure S6: Heterogeneity and specificity of telencephalic neuromodulatory networks

Figure S7: HCR-based visualization of marker genes for individual cell types

---

**Figure S1.**
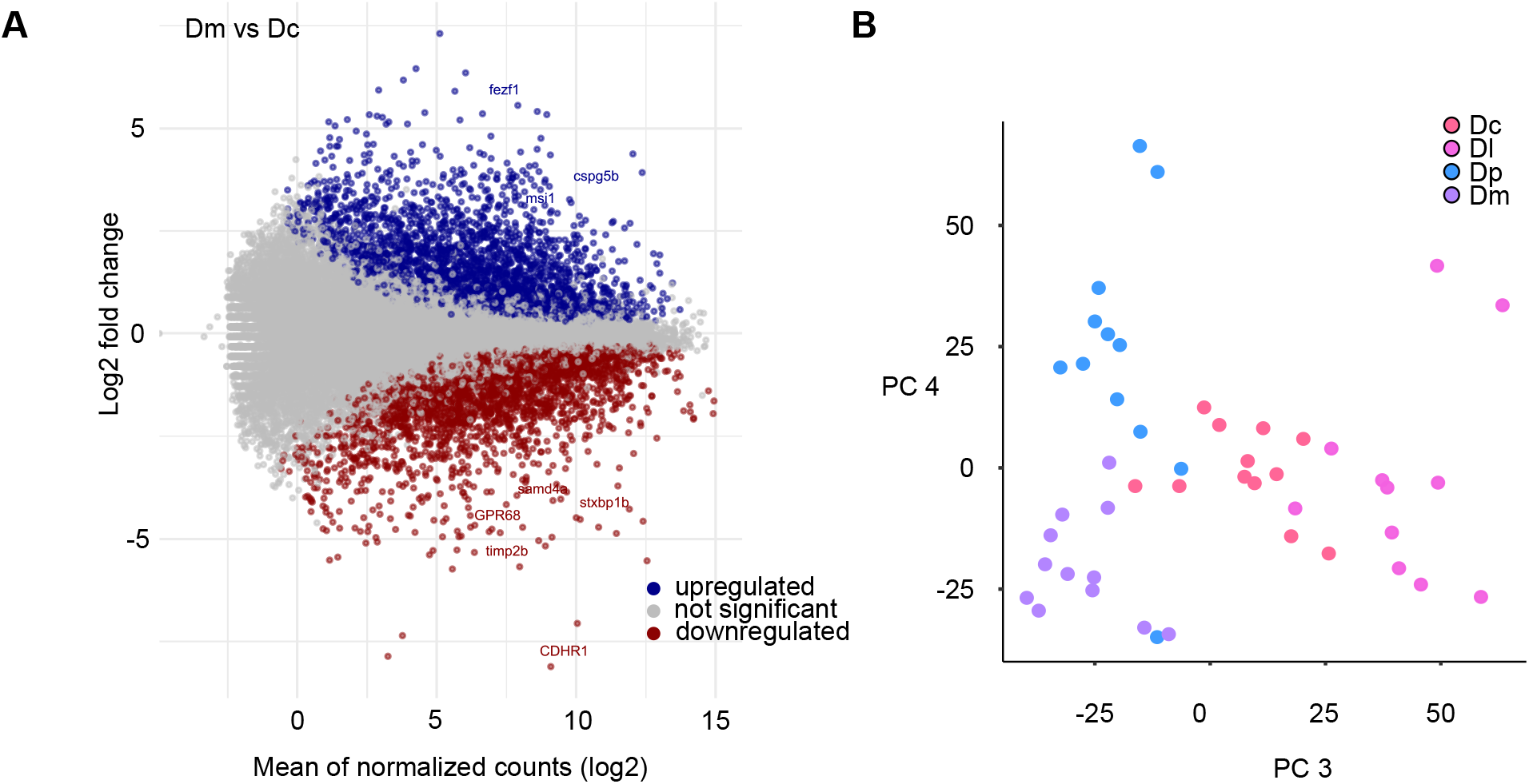
Bulk sequencing of pallial microdissected areas. (**A**) Example of differential gene expression in two different telencephalic brain areas (Dm and Dc). (**B**) Projection of gene expression patterns (2.000 most variable genes; same as in Fig. 1B) onto PCs 3 and 4. Each data point is one replicate from a pallial region (color-coded). Note separation of Dp and Dm. See Fig. 1B for projection of the same data onto PCs 1 and 2.

**Figure S2.**
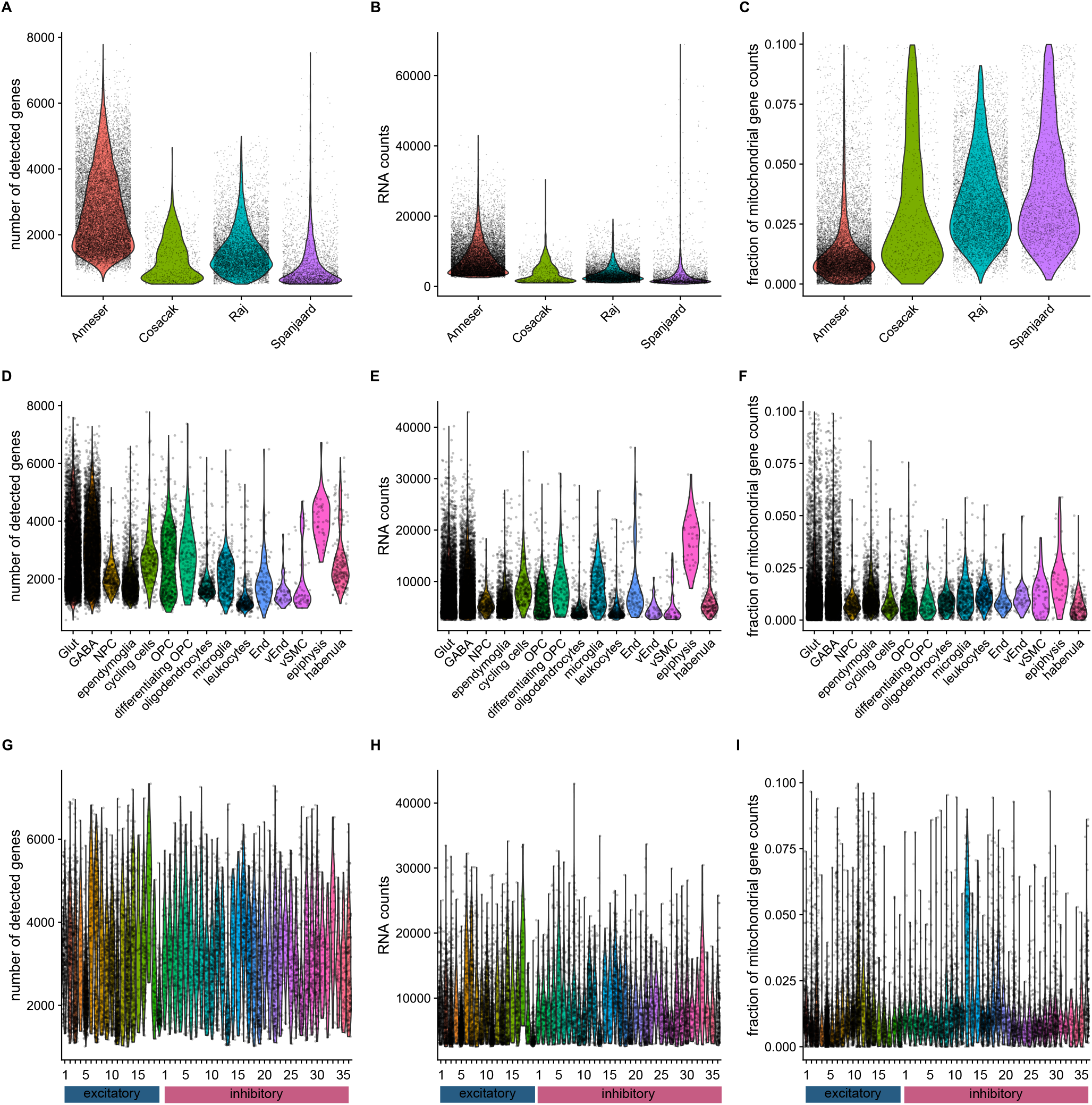
Quality assessment of scRNA-seq dataset. (**A-C**) Comparison of quality-related parameters of our single-cell RNA sequencing (scRNA-seq) dataset (“Anneser”) to previously published scRNA-seq datasets from the zebrafish telencephalon (“Cosacak”^89^, “Raj”^36^, “Spanjaard”^90^). Each dot represents one sequenced cell; violin plots show distributions. In our dataset, the number of detected genes and RNA counts are comparably high (**A**, **B**) while the fraction of mitochondrial gene reads, which usually indicates cell death, is relatively low (**C**). (**D-F**) Same quantifications stratified by superclusters in our dataset (Fig. 2A). (**E-I**) Same quantification stratified by neuronal subtypes.

**Figure S3.**
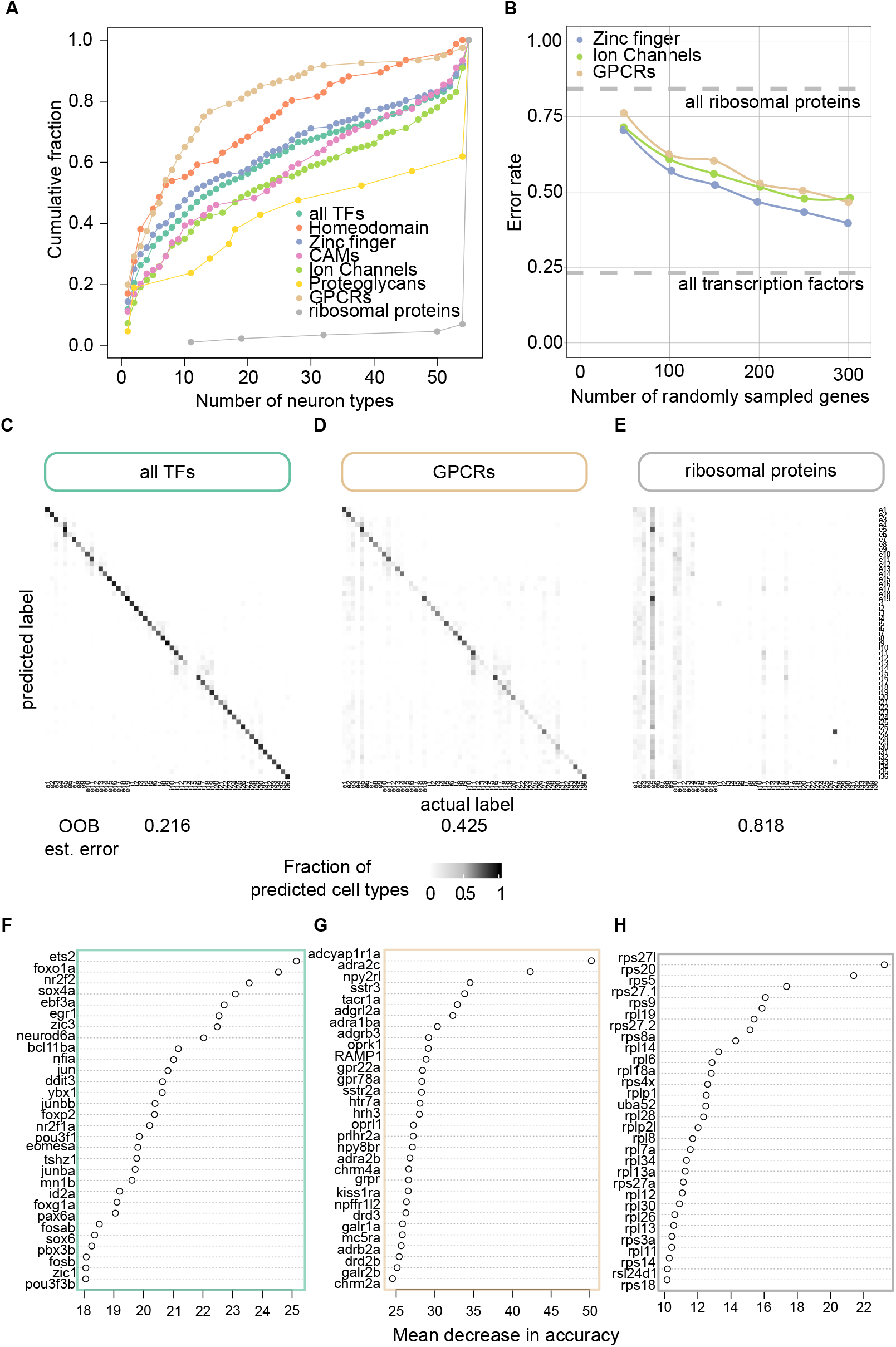
Cell-type specificity of gene expression. (**A**) Cumulative fraction of information about neuron types contained in expression patterns of different gene families (TFs, transcription factors; Homeodomain TFs; Zinc finger TFs, CAMs, cell adhesion molecules; ion channels; proteoglycans; ribosomal proteins). Information content of individual gene families was assessed by classifying the 55 neuronal cell types which were Louvain-clustered based on the entire transcriptome. This analysis is similar to Fig. 1D but follows the computation in Hain et al. 2022_27_. (**B**) Different accuracy in random forest classification might not only be attributable to degree of heterogeneity in the expression pattern, but simply to the number of genes available per gene family, as highlighted in this plot, in which we subsampled different gene families to the same size and plot against the accuracy of the corresponding random forest classifier, bounded by the accuracy of the ribosomal gene classifier and the classifier trained on the entire set of transcription factors. (**C**-**E**) Cell type classification performance of random forest classifiers trained on expression patterns of transcription factors, GPCRs, or ribosomal genes. The number below each plot indicates the out-of-bag error estimate. Note high performance of classifiers trained on TFs and GPCRs but not ribosomal genes. (**F-H**) Most informative genes for each of the classifier, ranked by mean decrease in accuracy when left out of the classifier.

**Figure S4.**
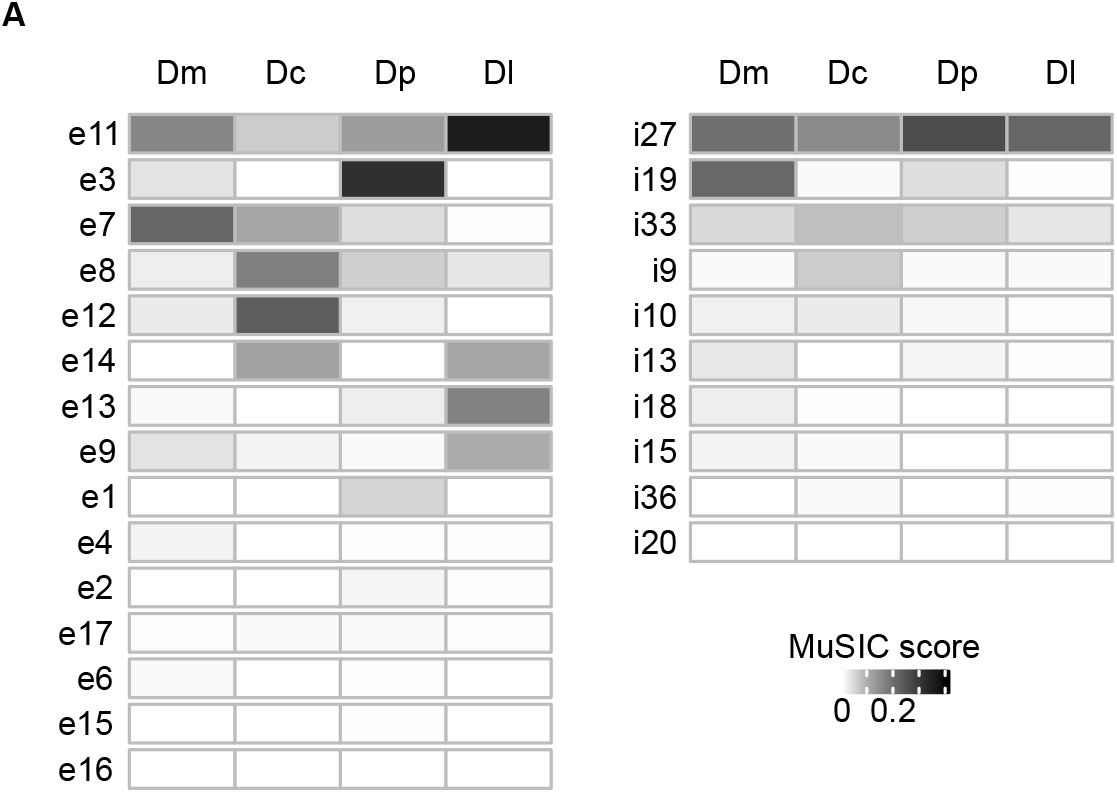
Contribution of individual cell types to pallial areas. (**A**) Deconvolved contribution of individual cell types to pallial areas, computed by the MuSIC algorithm^43^. Heatmaps are separated into glutamatergic and GABAergic cells and sorted by maximum contribution score.

**Figure S5.**
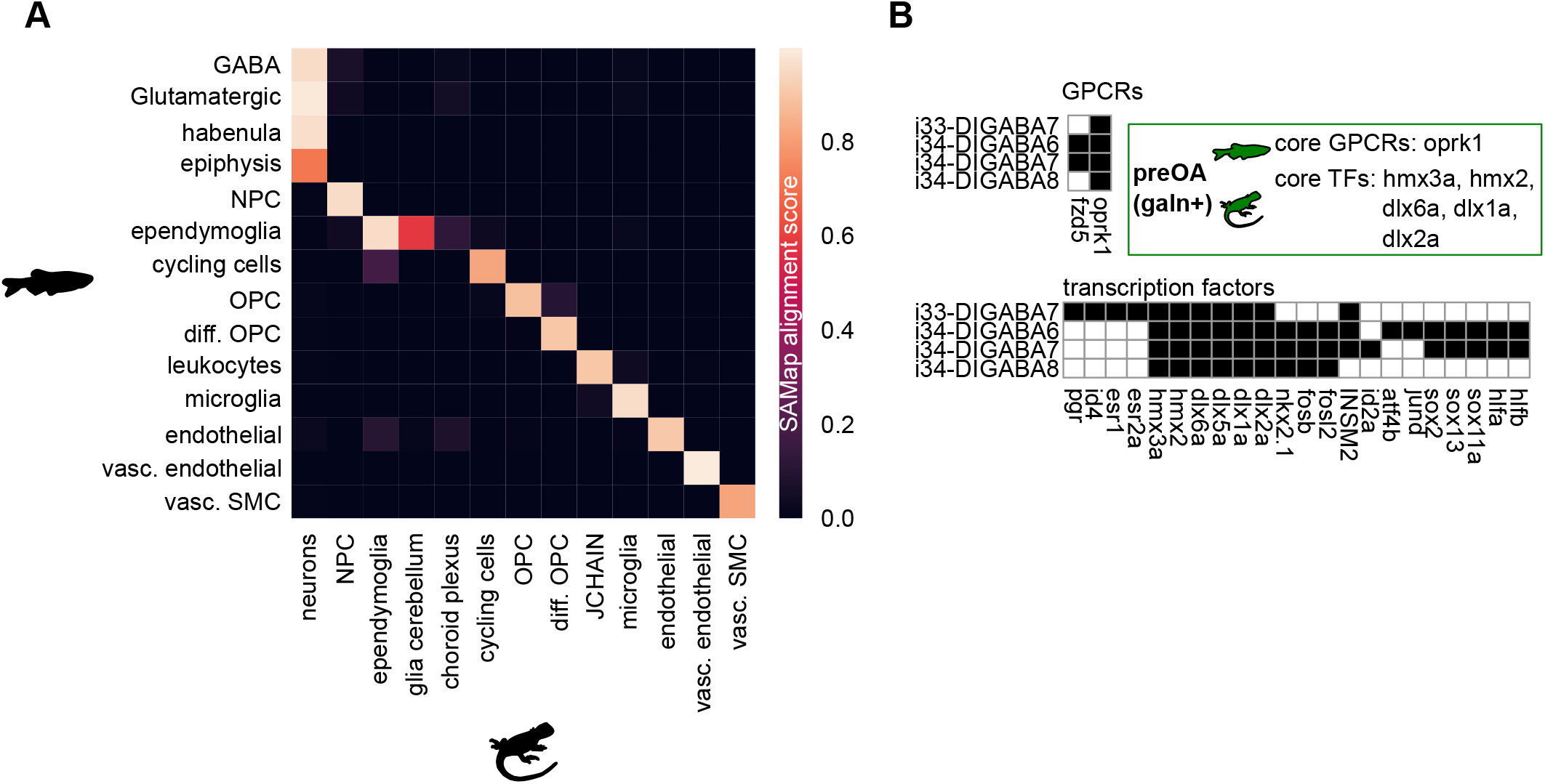
Evolutionary conservation of cell superclusters between zebrafish and bearded dragon. (**A**) Alignment score between corresponding superclusters in zebrafish and the bearded dragon. The sparsity of the matrix implies specificity of alignment. vasc. SMC, vascular smooth muscle cells; OPC, oligodendrocyte precursor cells; diff. OPC, differentiating OPC; NPC; neural progenitor cells, JCHAIN, plasma cells. (**B**) Binarized expression heatmap of GPCRs and TFs significantly contributing to the alignment of galanin-expressing cells in the preoptic area of the zebrafish and hypothalamic galanin+ cells in the lizard. Both appear to be susceptible to modulation via the K-opioid receptor in both species.

**Figure S6.**
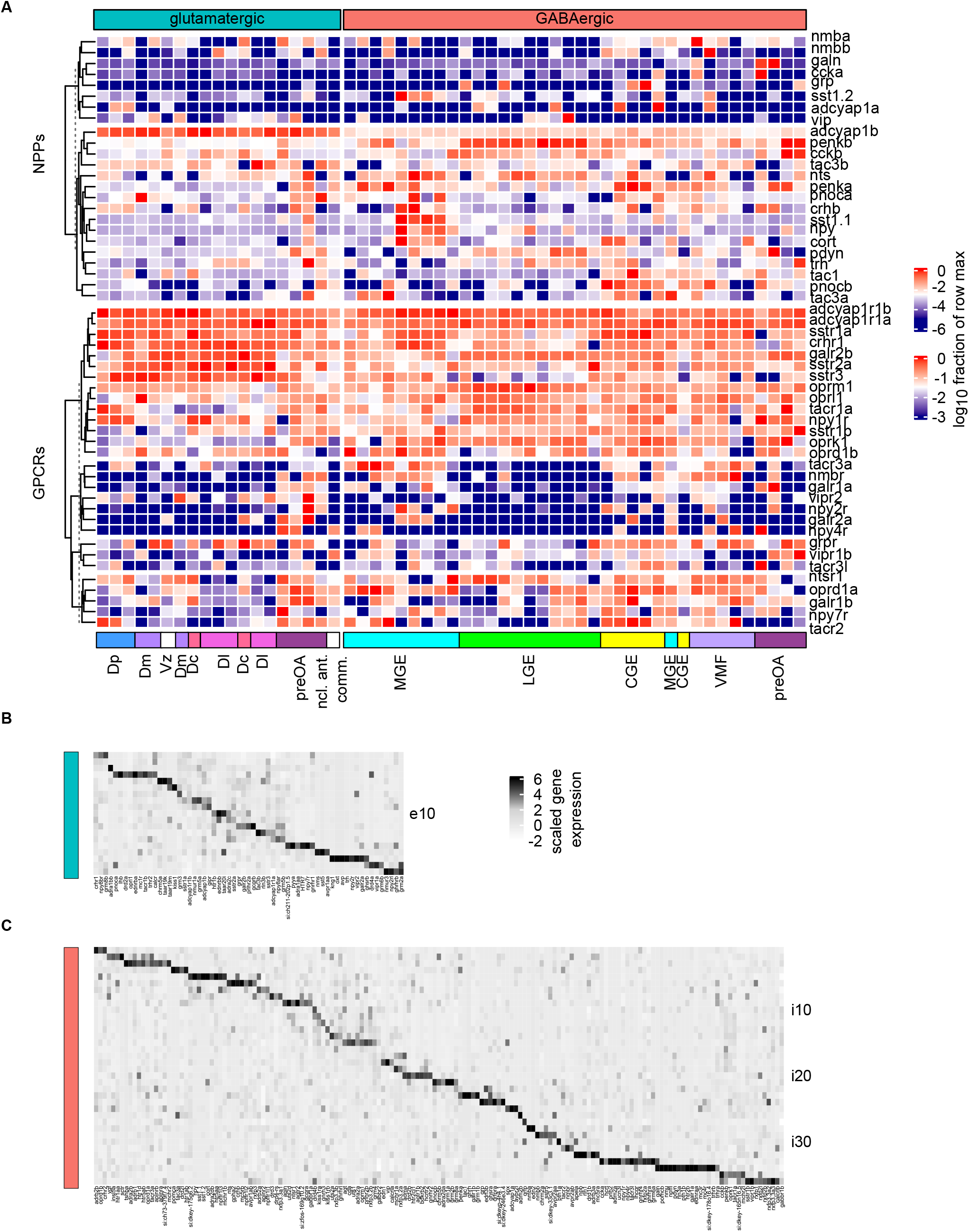
Heterogeneity and specificity of telencephalic neuromodulatory networks. (**A**) Normalized expression of selected NPPs and GPCRs in GABAergic and glutamatergic cell types identified in our dataset. Expression is shown using a logarithmic color scale after normalization to the maximum in each row. Rows were ordered by hierarchical clustering (left). (**B, C**) Enlargements of Fig. 6J showing normalized (z-scored) expression levels of all genes associated with neuromodulation separately in glutamatergic and GABAergic cell types. Matrices were sorted by maximum expression per cell type.

**Figure S7.**
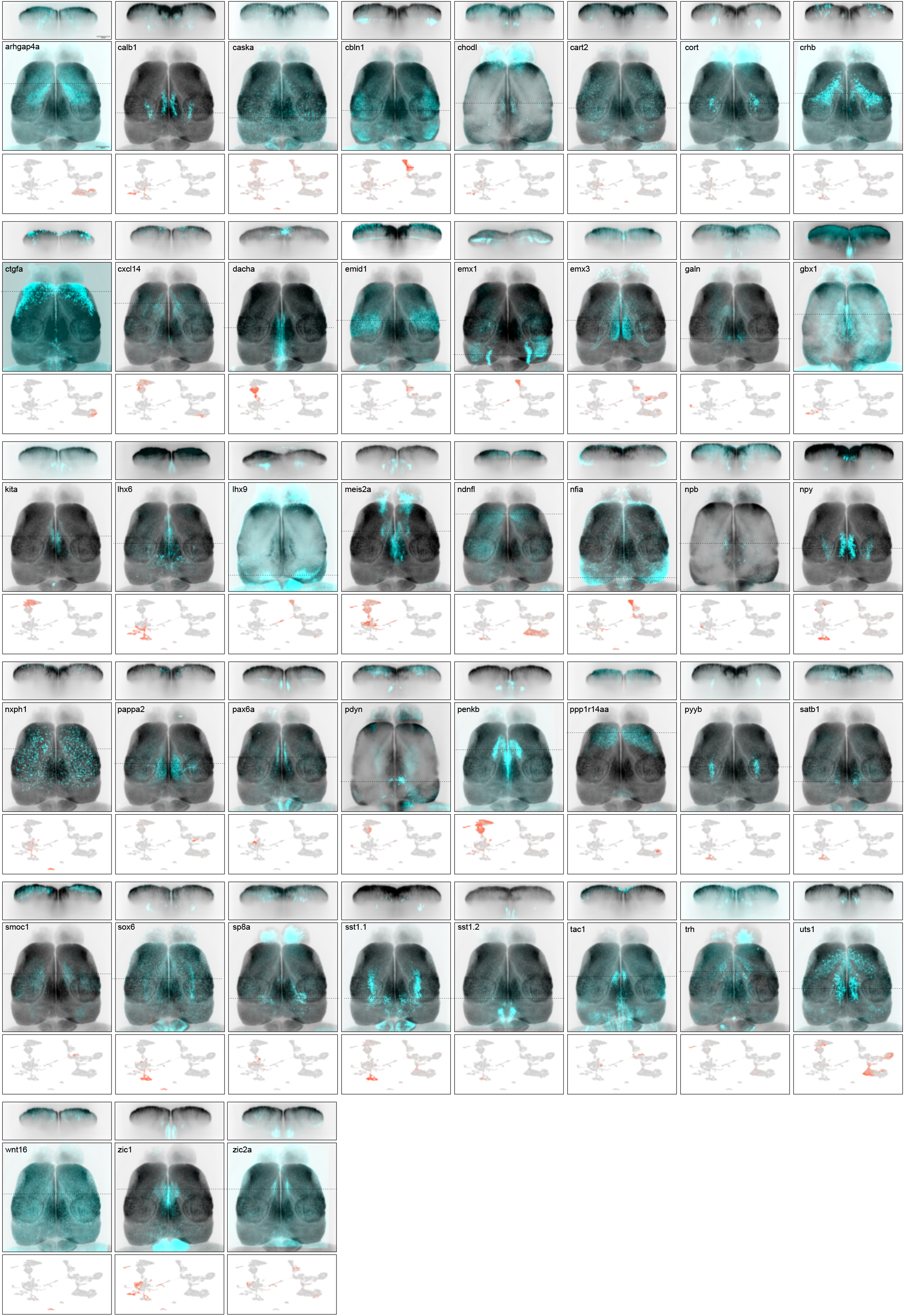
HCR-based visualization of marker genes for individual cell types. Expression patterns of 43 marker genes in the adult telencephalon. Images show maximum-intensity z-projections (dorsal views) and a coronal section at the anterior-posterior position indicated by the dashed line. Expression patterns were determined by HCR (cyan) and mapped onto a common reference brain using *snap25a* expression for registration (gray background). Scale bar: approximately 200 μm (not precise due to registration of image datasets onto the reference brain). Plots below show gene expression levels in individual cells mapped onto the first two UMAP components (Fig. 2C). Genes are ordered alphabetically.

**S2.**
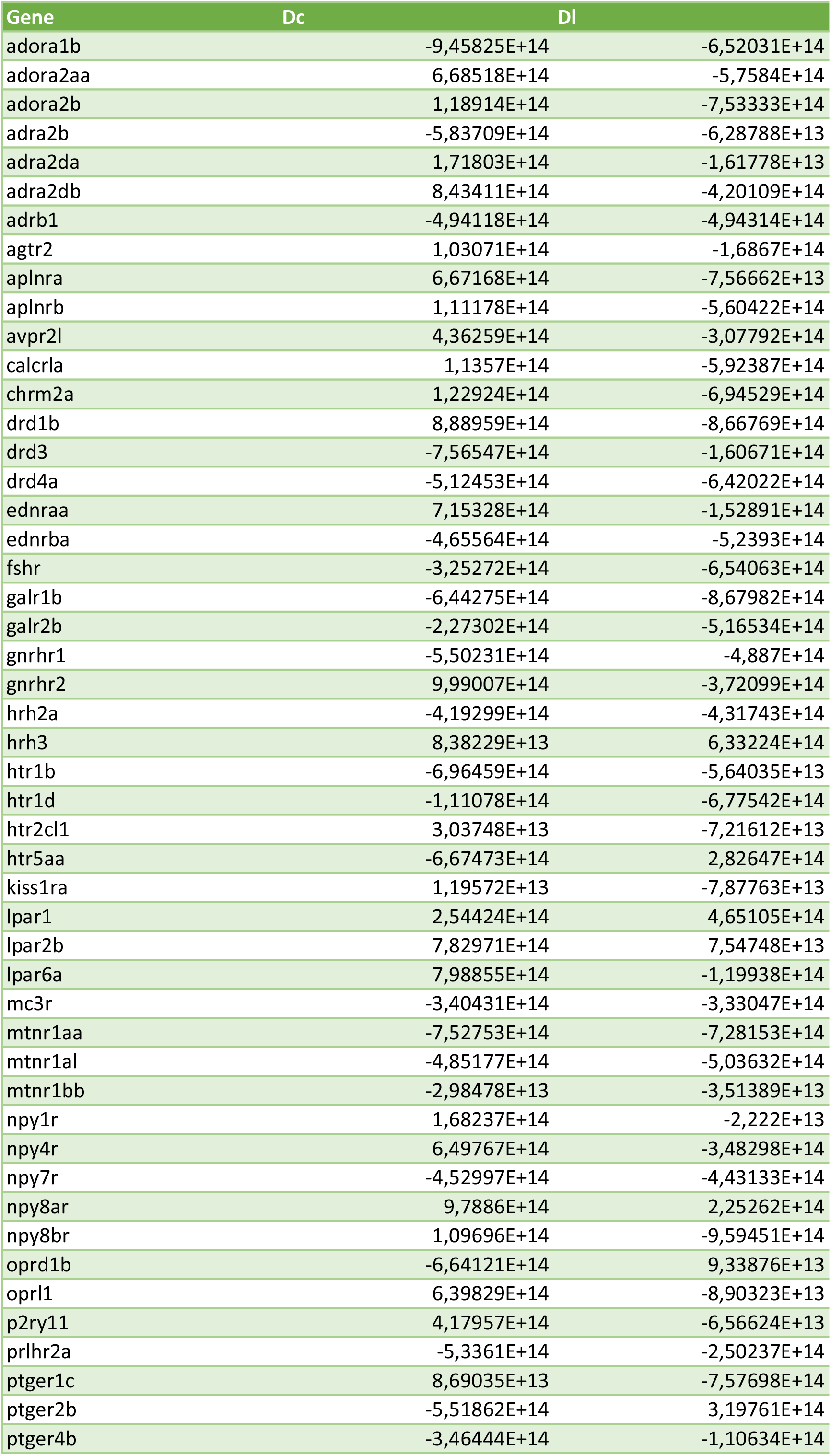
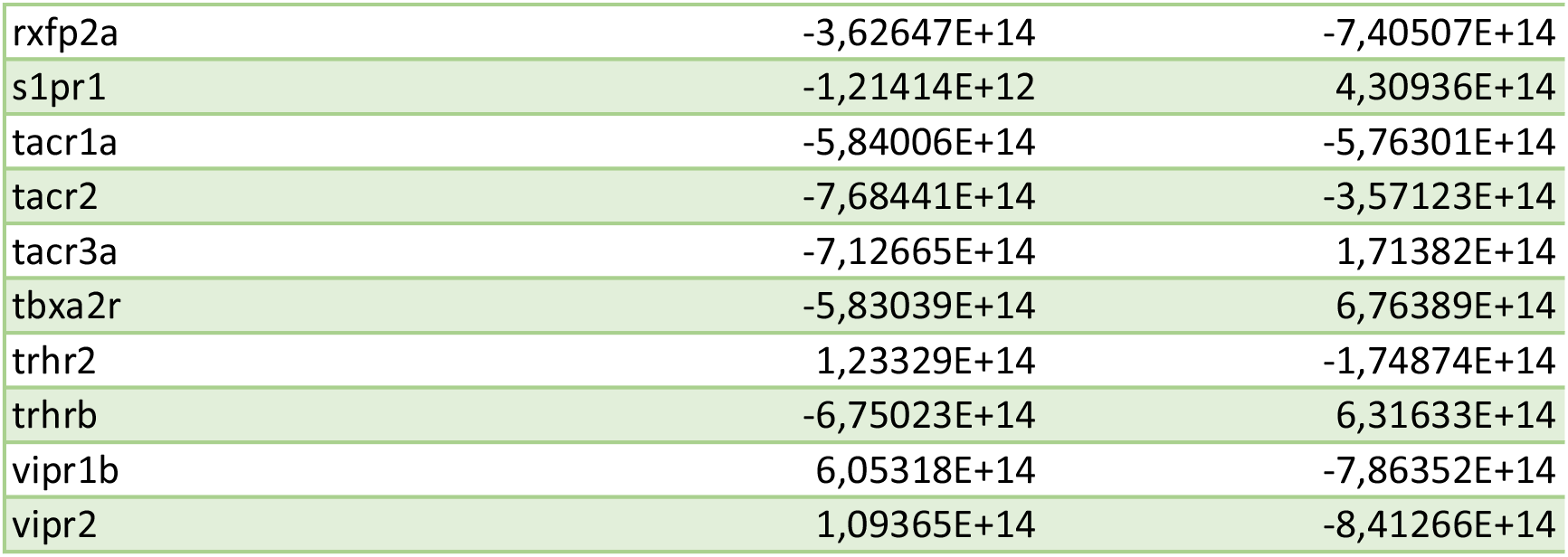
List of GPCRs and their normalized bulk expression.

| Dp | Dm |
| --- | --- |
| 7,99502E+14 | 7,44018E+14 |
| -2,39793E+14 | 9,91287E+14 |
| -4,50568E+14 | -4,80202E+14 |
| 6,21581E+14 | 5,38516E+14 |
| 8,16875E+14 | -8,40383E+14 |
| 3,44933E+14 | -4,6176E+14 |
| -3,64624E+14 | 1,31186E+14 |
| -1,59867E+14 | -7,16228E+14 |
| -1,34282E+13 | -4,63526E+14 |
| -2,29096E+13 | -3,68961E+14 |
| 2,93671E+14 | -4,47788E+14 |
| -7,10123E+14 | 1,17449E+14 |
| -3,87284E+14 | -5,53857E+13 |
| -3,40526E+14 | -4,68979E+14 |
| 6,5099E+14 | 2,52839E+14 |
| -1,28113E+14 | 1,22909E+14 |
| 1,66488E+14 | -7,41666E+14 |
| 1,27345E+14 | 8,18488E+14 |
| -1,47765E+14 | 1,07259E+14 |
| 3,49302E+14 | 1,09062E+14 |
| -5,21563E+13 | 1,22236E+14 |
| 8,32982E+14 | 1,65224E+14 |
| -3,29622E+14 | -3,28293E+13 |
| -3,52709E+14 | 1,16777E+13 |
| -8,05994E+14 | -6,12691E+14 |
| -9,91826E+14 | 1,31267E+14 |
| -3,70167E+13 | 1,10233E+14 |
| 8,28483E+14 | -4,70753E+14 |
| -2,70478E+13 | 6,78858E+13 |
| -6,60884E+14 | 1,87281E+13 |
| 2,35521E+14 | -9,16292E+14 |
| -1,58815E+14 | -6,93342E+14 |
| -9,69255E+14 | -5,91987E+14 |
| -1,55527E+13 | 6,61277E+14 |
| 4,53833E+14 | 9,66394E+13 |
| -3,7992E+14 | 1,32676E+14 |
| -1,40301E+14 | 7,60886E+14 |
| 6,62245E+14 | -6,26799E+14 |
| 7,44631E+14 | -4,04957E+14 |
| -1,76072E+14 | 1,03527E+14 |
| -7,44082E+14 | -4,41269E+14 |
| 2,93271E+14 | -5,10739E+14 |
| -7,05393E+14 | 5,13461E+14 |
| -5,23955E+14 | -1,15058E+14 |
| 3,96376E+14 | -3,6173E+14 |
| 9,71967E+14 | 7,62022E+14 |
| 5,73764E+14 | -7,48242E+14 |
| -4,15572E+13 | 9,38122E+14 |
| -3,94057E+14 | 8,41916E+14 |
| -3,19513E+14 | 1,36096E+14 |
| 3,39811E+14 | 7,85134E+14 |
| 1,01437E+14 | 9,7907E+14 |
| 8,30467E+14 | 2,65336E+14 |
| 4,1487E+14 | 1,40696E+14 |
| -5,14873E+14 | 1,03591E+14 |
| -5,08123E+14 | -5,64869E+14 |
| -2,80313E+14 | 8,97436E+14 |
| 5,56819E+14 | -4,41315E+14 |
| -5,71949E+14 | -2,65294E+14 |

